# *B. pertussis* tracheal cytotoxin biases NOD signaling to suppress IL-1 mediated inflammation and evade adaptive immunity

**DOI:** 10.1101/2025.08.08.669383

**Authors:** David M. Rickert, Elizabeth M. Hill, Bella Carnahan, Sasha Cardozo, Riley E. Himmelberger, Neeta Rajbanshi, Tialfi Bergamin de Castro, Ravi Bharadwaj, Katelyn M. Parrish, Nicholas J. First, Monica C. Gestal, Alison J. Scott, Neal Silverman, William E. Goldman, Nicholas H. Carbonetti, Karen M. Scanlon, Catherine Leimkuhler Grimes, Ciaran Skerry

## Abstract

*Bordetella pertussis* releases the monomeric peptidoglycan (PGN) fragment tracheal cytotoxin (TCT) due to inefficient recycling by the permease AmpG. Releasing this PGN is metabolically costly and potentially immune alarming and the benefits to *B. pertussis* are unclear. While TCT has been characterized as a potent NOD1 agonist capable of causing the extrusion of ciliated cells, in vitro, the consequences of its release have yet to be studied *in vivo*. Here we show that selective PGN release by *B. pertussis* biases host PGN sensing toward NOD1 and away from NOD2, suppressing IL-1β-driven inflammation and blunting adaptive immune recruitment. Mice infected with a TCT over-releasing strain (TCT(+)) exhibit reduced pulmonary immunopathology relative to wild type (WT) and a TCT-under-releasing strain (TCT(-)), despite similar bacterial burdens. NOD reporter assays demonstrate that TCT release enhances NOD1 activation and inversely correlates with NOD2 activation. Bulk transcriptomic analysis of infected lungs shows that *B. pertussis* PGN release dampens pro-inflammatory transcriptional programs. Single-cell transcriptomic determined *Nod2* expression is limited to inflammatory myeloid subsets. IL-1 family genes were highly enriched in *Nod2-* but not *Nod1* expressing alveolar macrophages. Upstream regulator analysis predicted IL-1β as a major driver of *B. pertussis* inflammation, which was enhanced by the absence of PGN release. Flow cytometry shows that PGN release skews macrophages polarization toward M2 and away from M1 in a NOD1 dependent manner. Finally, extracellular release of PGN and subsequent reduced IL-1 production facilitated the suppression fibroblast chemokine programs (e.g., CXCL13, CCL19), diminished recruitment of B and T cells, reduced iBALT formation, and limited immune memory development. Conversely, IL-1R1 deficiency impairs adaptive recruitment and bacterial clearance despite similar innate infiltration. Together, these data suggest PGN release by *B. pertussis* is an immune-evasion strategy, favoring NOD1 activation over NOD2, reducing IL-1–dependent fibroblast reprogramming, and curtailing chemokine-driven adaptive responses.

**Figure 8.**
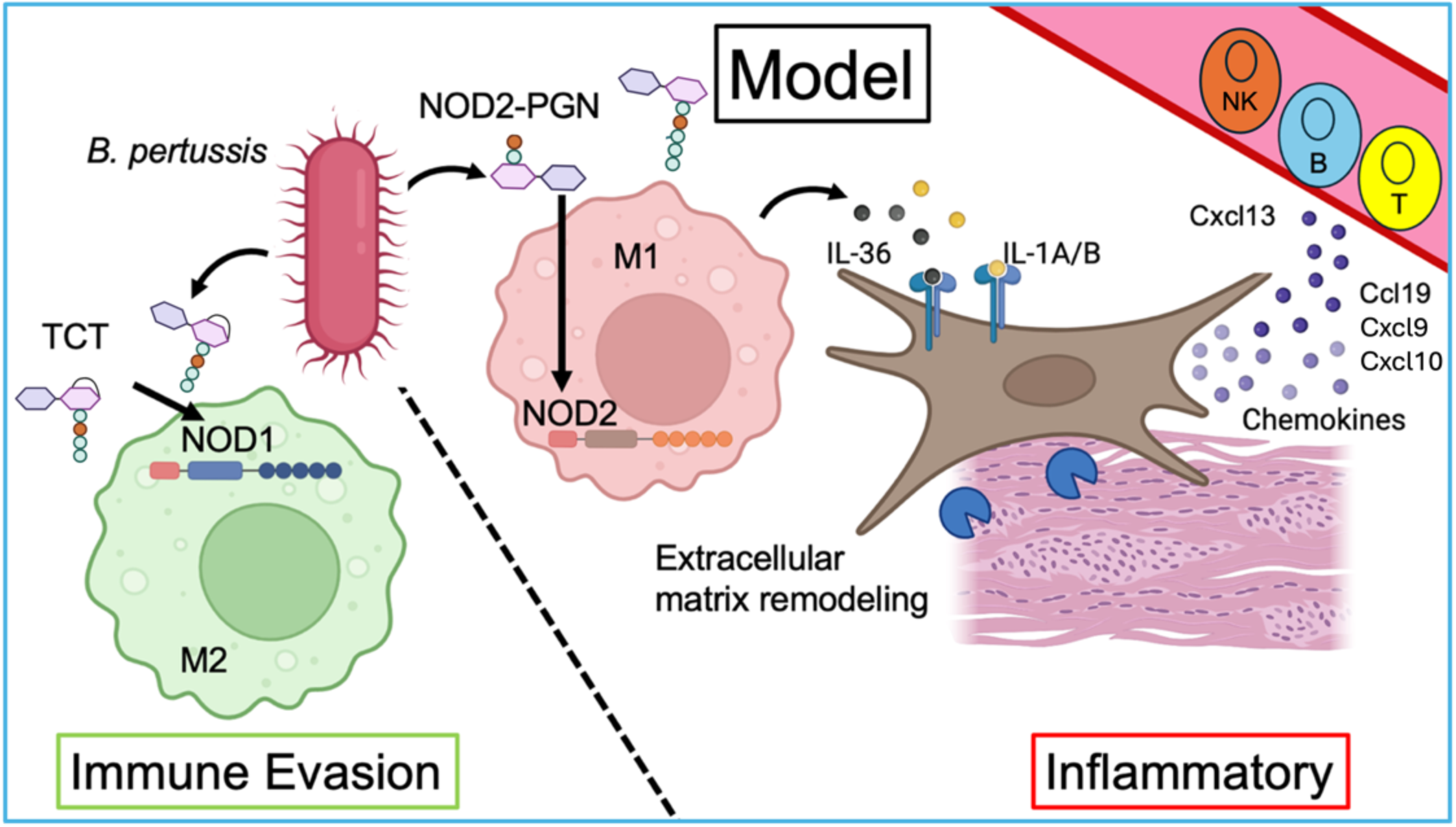
Graphic Abstract *B. pertussis* can produce both NOD1 and NOD2 activating PGNs. Release of TCT promotes NOD1 activation and diminishes NOD2 activation. NOD2 activation in myeloid cells drives M1 polarization of macrophages and IL-1 family cytokine production. IL-1 family cytokines skew fibroblasts towards an inflammatory phenotype, leading to chemokine release, extracellular remodeling, and recruitment of lymphocytes. Therefore, TCT release tempers long-term immunity to *B. pertussis*.

## INTRODUCTION

Bacterial pathogens have evolved diverse mechanisms to manipulate host responses and promote their persistence. Many pathogens strategically alter the structure or availability of pathogen-associated molecular patterns (PAMPs) to shape the host immune responses. For example, modifications in the structure of the lipid A region of lipopolysaccharide (LPS) determine the magnitude of the host immune responses^1^. However, the impact of peptidoglycan (PGN) structure on host responses to bacterial infections is under appreciated.

Peptidoglycan, a conserved component of the bacterial cell wall, forms a mesh-like polymer of repeating disaccharides crosslinked by short peptides^2^. Cytosolic receptors NOD1 and NOD2 detect PGN motifs; such as Diaminopimelic acid (DAP) and muramyl dipeptide (MDP) or NagK phosphorylated MDP, triggering NF-kB-mediated inflammatory signaling^3–5^. This places selective pressure on bacterial pathogens to modify muropeptide release or the biochemical structure to evade immune responses.

Interestingly, several unrelated bacterial pathogens have independently evolved the ability to actively release specific muropeptides during infection. *Borrelia burgdorferi*, *Neisseria gonorrheae* and *Bordetella pertussis* release free PGN fragments, an energetically costly process, which could increase immune detection^6,7^. Both *N. gonorrheae* and *B. pertussis* preferentially release a GlcNAc(beta1-4)-MurNAc(1,6-anhydro) DAP-type tetrapeptide, termed TCT in *B. pertussis*^8,9^. Excitingly, *B. pertussis* and *N. gonorrheae* evolved divergent mechanism to modulate TCT release that converge through the same player, bacterial permease AmpG. Typically, AmpG recycles anhydro-muropeptides from the periplasm into the cytoplasm for reuse^10^. *B. pertussis* evolved an insertion sequence in the promoter of its endogenous *ampG*, decreasing expression^11^. On the other hand, *N. gonorrheae* hamstrung its AmpG by evolving mutations which decreased efficiency of transport^9^. These evolutionary observations prompt the question of why these diverse pathogens would impair their AmpG permeases to promote the release of extracellular PGN fragments like TCT.

*In vitro* studies have demonstrated that TCT promotes extrusion of ciliated cells, thereby impairing mucociliary clearance of the pathogen^12^. Additional *in vitro* studies have linked TCT to iNOS expression, IL-6 production and break down of barrier integrity^13^. To assess the benefit of reduced AmpG permease efficiency *in vivo*, we investigated the impact of TCT in *B. pertussis* pathology and immunity. We employed *B. pertussis* strains engineered to overproduce or limit TCT to assess its role *in vivo*. *B. pertussis* expresses an inefficient AmpG, resulting in periplasmic accumulation and extracellular release of the disaccharide-tetrapeptide TCT^6,14^. Replacing the native *ampG* with the more efficient *E. coli* version reduced TCT release by ∼98% (TCT(-)) without affecting bacterial growth^15,11^. Conversely, deletion of the native *B. pertussis ampG* resulted in a 24-fold increase in TCT release without compromising growth, producing a TCT-over-releasing strain (TCT(+))^11^.

To evaluate the *in vivo* impact of TCT release, we challenged mice with TCT(+), TCT(-) or wild-type (WT) strains of *B. pertussis*. Contrary to prior *in vitro* studies linking TCT to inflammation, we observed an unexpected anti-inflammatory effect of TCT *in vivo*. Reporter assays and single cell transcriptomics revealed that TCT selectively engages NOD1 over NOD2 contributing to its anti-inflammatory effects. We identify divergent outcomes of signaling via NOD1 and NOD2, highlighting a NOD2-IL1 signaling axis in which IL-1 activates IL1R1+ airway fibroblasts to promote inflammatory cytokine and chemokine production and extracellular matrix remodeling. Our data suggests *B. pertussis* exploits TCT release to bias host sensing of PGN towards NOD1, thereby avoiding pro-inflammatory NOD2 activation. This TCT-mediated immune skewing impairs adaptive immune recruitment and diminishes effective immune memory responses. Further, it highlights divergent outcomes of PGN signaling via NOD1 or NOD2 during infection. These findings have broad implications for understanding bacterial immune evasion, PGN sensing, and vaccine adjuvant development.

## RESULTS

### TCT dampens pulmonary inflammation following *B. pertussis* challenge

To assess the potential benefits of PGN release, we investigated the contribution of tracheal cytotoxin (TCT) to the pathogenesis of *Bordetella pertussis* infection *in vivo*. We intranasally challenged mice with WT, TCT(+), or TCT(-) strains of *B. pertussis*. Based on prior *in vitro* studies describing TCT as a pro-inflammatory PAMP, we hypothesized that infection with TCT(+) *B. pertussis* would induce heightened pulmonary inflammation compared to WT or TCT(-). Surprisingly, we observed the opposite outcome: mice infected with TCT(+) exhibited significantly reduced lung inflammation compared to those infected with the WT strain (Figure 1A–D). Histological analysis of lungs at 7 days post-infection (dpi) revealed diminished immune cell infiltration and less bronchovascular bundle cuffing following infection with TCT(+) than WT or TCT(-) (Figure 1A–C). To determine whether these differences in immunopathology were due to discrepancies in bacterial load, we quantified lung colony-forming units (CFUs) at 2 and 7-dpi. No significant differences in CFU were observed between strains at either time point (Figure 1E), indicating that inflammation was not driven by discrepancies in bacterial burden. We also observed comparable growth kinetics between strains *in vitro* in Stainer-Scholte medium, further suggesting differences in pathology were not due to fitness defects in the mutant strains (Sup. Figure 1A, B).

**Figure 1.**
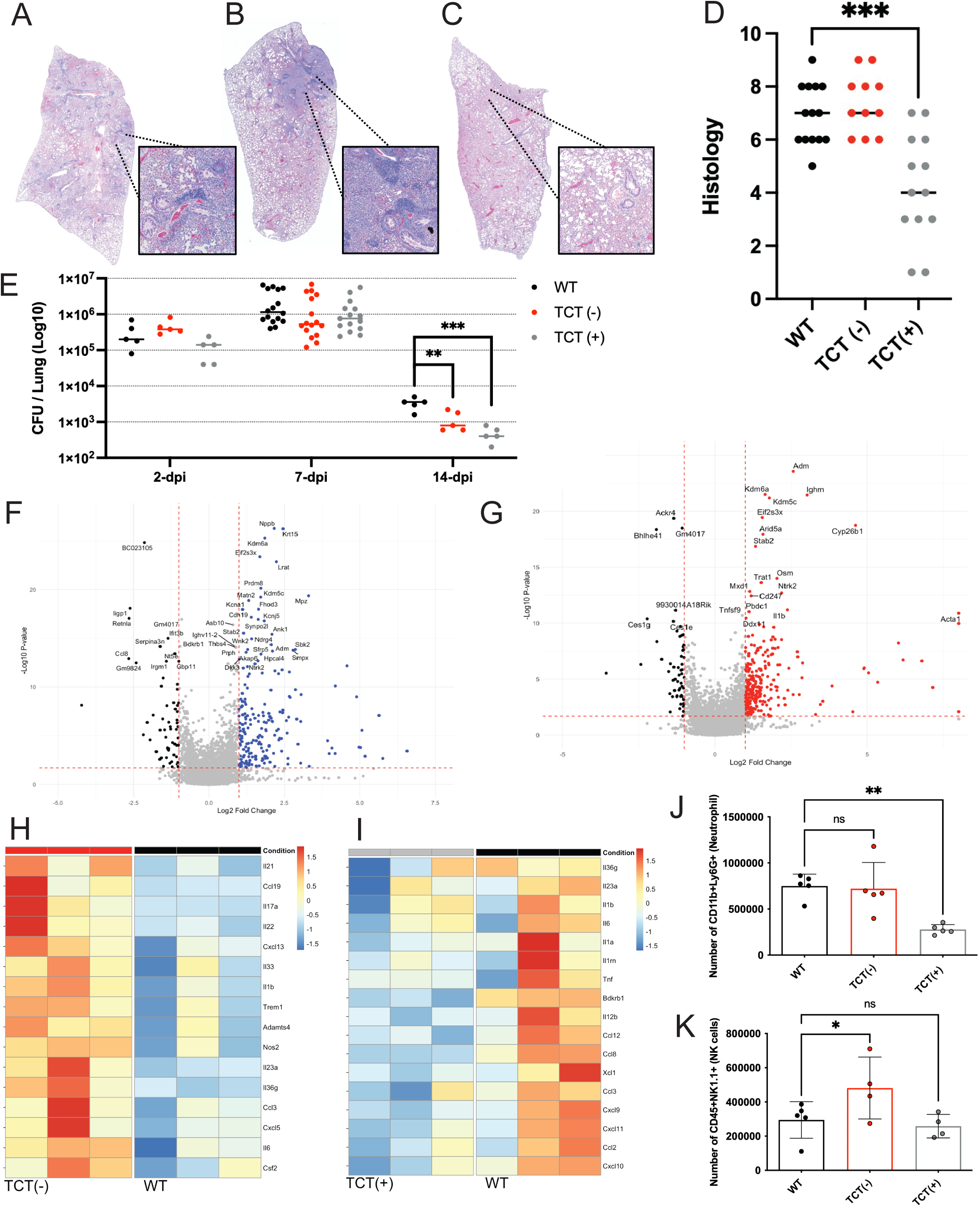
Representative 40x stitched H&E stains of *B. pertussis* strains (**A**) WT, (**B**) TCT(-), and (**C**) TCT(+) 7-dpi BALB/c mouse lungs and **(D)** inflammatory histology scores for mice intranasally challenged with 2x10^6^ CFU. (**E**) Bacterial burden in the lungs at 2, 7, and 14-dpi. Bulk transcriptomic volcano plots comparing 4-dpi lungs of BALB/c mice infected with (**F**) WT vs TCT(-) or (**G**) WT vs TCT(+). Heat map of bulk transcriptomic comparisons, highlighting cytokines and chemokines which were differentially regulated between (**H**) WT vs TCT(-) or (**I**) WT vs TCT(+). (**J**) Number of CD45+CD11b+Ly6g+ (Neutrophils) cells in the 7-dpi lungs of BALB/c mice challenged with WT, TCT(-), or TCT(+) strains of *B. pertussis*. (**K**) Number of CD45+NK1.1+ (NK) cells in the 4-dpi lungs of BALB/c mice challenged with WT, TCT(-), or TCT(+) strains of *B. pertussis*.

These findings suggest that TCT release dampens inflammatory signaling during infection in the lung leading the divergent pulmonary inflammation observed at 7-dpi. To further assess the nature of this immune-dampening, we performed bulk RNA sequencing of 4-dpi lungs from mice infected with WT, TCT(-), or TCT(+) strains. Consistent with earlier histopathology data, mice infected with the TCT(-) strain exhibited a pronounced upregulation of pro-inflammatory cytokine transcripts, whereas TCT(+)-infected mice had reduced transcription of inflammatory cytokines compared to WT (Figure 1F, H). Notably, TCT(-) infection induced significantly higher expression of inflammatory cytokines and chemokines such al *Il17a*, *Il21, Il23a, Il36a, Ccl19*, *Cxcl5*, and *Cxcl13* (Figure 1F, H).

Canonically, cytokines *Il21* and *Il23a* play critical roles in skewing towards Th17 immunity and production of Il17a^16,17^. Il17 signaling results in increased expression of chemokines, e.g., Cxcl5, to recruit neutrophils to the site of the infection and is critical for immunity to *B. pertussis*^18,19^. The upregulation of these cytokines and chemokines in the absence of TCT release by *B. pertussis* suggests TCT dampens this pathway to inhibit neutrophil recruitment. To test this hypothesis, we performed flow cytometry to assess the numbers of neutrophils recruited to the lung during infection by different TCT strains. Supporting our hypothesis, we discovered that neutrophil recruitment to the lung was impacted by TCT release (Figure 1J). Mice infected with TCT-over releasing strain demonstrated significantly decreased neutrophil numbers in the lungs, highlighting one mechanism by which TCT may suppress pulmonary inflammation (Figure 1J).

Conversely, TCT(+) infection led to significantly reduced expression of cytokines and chemokines like *Il12b, Xcl1, Ccl8, Ccl12, Ccl19, Cxcl9, Cxcl10* and *Cxcl11* relative to WT infection (Figure 1G, I). Chemokines like Cxcl9, Cxcl10, and Cxcl11 are critical for recruiting lymphocytes like NK cells to sites of injury, infection, and inflammation^20^. NK cells play critical roles in *B. pertussis* immunity via release of IFN-𝛾^21,22^. Given that increased TCT release dampened the production of these chemokine transcripts, we hypothesized that TCT suppresses recruitment of NK cells. Flow cytometry analysis further supported this, as mice infected with TCT-under releasing strains of *B. pertussis* demonstrated increased NK cells in the lung (Figure 1K), implicating another cellular mechanism by which TCT repressed inflammation in the lung, and benefits the pathogen by delaying recruitment of key mediators of immunity to *B. pertussis* infection.

Interestingly, *il1b* and *nos2*, genes associated with M1 macrophage polarization, were upregulated in TCT(-) infected lungs, suggesting a link between TCT release and the inhibition of M1 polarization (Figure 1F, H). Contrary to previous *in vitro* data, these studies demonstrate that TCT release may facilitate the evasion of host responses by dampening inflammatory responses, further, we suggest this may be due to the skewing of macrophage polarization toward M2 phenotypes and suppression of key pro-inflammatory pathways.

### Increased release of TCT by *B. pertussis* prevents the release of NOD2-stimulatory muropeptides

We next sought to determine the mechanisms by which TCT dampens inflammatory immune responses. TCT is known to mediate immune signaling via NOD1^23^. We confirmed this by treating human and murine NOD1 and NOD2 reporter cells with purified TCT, and validated that TCT is a potent NOD1 agonist, with a preference for murine NOD1 (Sup. Figure 2A, B). TCT did not stimulate either murine or human NOD2 response (Sup. Figure 2C, D). To gain further insight into TCT-mediated skewing of immune responses and determine if TCT release by *B. pertussis* altered activation of NOD1 and NOD2, we co-incubated WT, TCT(-) and TCT(+) strains with murine NOD1 and NOD2 reporter cells. As expected, improving AmpG permease efficiency and impairing TCT release diminished NOD1 responses, while reduced permease efficiency increased TCT release and enhanced NOD1 signaling (Figure 2A). However, surprisingly, the opposite pattern was observed for NOD2 signaling; greater TCT release generated a weaker NOD2 response as diminished TCT release enhanced NOD2 signaling (Figure 2B). These data demonstrate that *B. pertussis* PGN can activate NOD2 and that greater TCT release inversely decreases NOD2 sensing (Figure 2A, B). This could potentially be due to masking or competitively inhibiting access to NOD2-stimulatory muropeptides.

**Figure 2.**
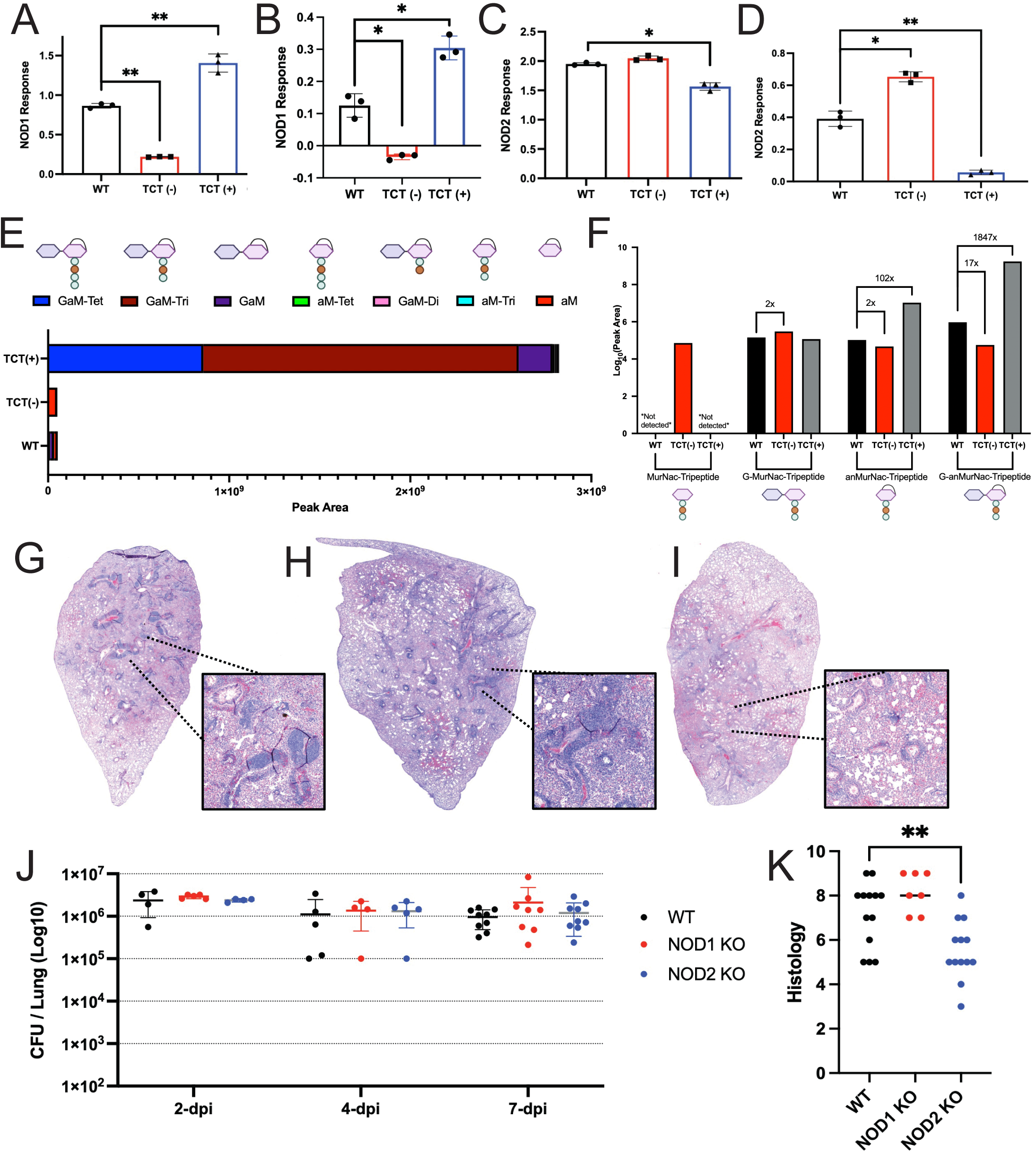
Murine NOD1 response to either (**A**) live *B. pertussis* (MOI 400) or (**B**) *B. pertussis* conditioned media (5%) and measured by HEK293 reporter cells with an NF-kB based SEAP reporter, assessed at 655nm. Murine NOD1 response to either (**C**) live *B. pertussis* (MOI 400) or (**D**) *B. pertussis* conditioned media (5%) and measured by HEK293 reporter cells with an NF-kB based SEAP reporter, assessed at 655nm. (**E**) Relative abundance of PGN fragments found in the conditioned media of WT, TCT(-), and TCT(+) strains of *B. pertussis*, measured by LC-MS, visualized by peak area. (**F**) Relative abundance of muro-tripeptides found in the conditioned media of WT, TCT(-), and TCT(+) strains of *B. pertussis*, measured by LC-MS, visualized by peak area. H&E stains of C57/B6 mice intranasally challenged with 2x10^6^ CFU for (**G**) WT, (**H**) NOD1 KO and (**I**) NOD2 KO 7-dpi lungs, (**J**) bacterial burden in the lungs at 2, 4, and 7-dpi, (**K**) inflammatory lung histology scores at 7-dpi.

To determine if the NOD activation profiles observed were due to secreted bacterial products like TCT, we evaluated the NOD activating profiles of conditioned media collected from the WT, TCT(-), and TCT(+) strains. Excitingly, these experiments mirrored the findings from whole-cell co-incubations.

Conditioned media from the TCT(+) strain induced stronger NOD1 but weaker NOD2 activation, while media from the TCT(-) strain elicited stronger NOD2 but weaker NOD1 activation (Figure 2C, D). These data support the hypothesis that TCT is the dominant NOD1 agonist in the media. But more excitingly, these data suggest that greater release of TCT leads to decreased release or presence of NOD2-stimulatory PGN fragments. Thus, the efficiency of AmpG, the permease mediating the differences in TCT release, may determine the muropeptide structures released extracellularly and activate either NOD1 or NOD2. We hypothesized that the increased NOD2 signaling observed in TCT(-) infection, may result from distinct, unidentified NOD2 agonist(s) that are otherwise retained or masked in the presence of inefficient PGN processing by AmpG and TCT release.

To investigate the impact of AmpG efficiency on the structures of extracellular muropeptides we performed high resolution liquid chromatography mass spectrometry to identify, and quantify the specific muropeptide species present in the WT, TCT(-), or TCT(+) conditioned media. First, we confirmed the impact of AmpG efficiency on TCT release. We measured 84-fold greater release of TCT by the TCT(+) strain compared to the WT parental strain (Sup. Figure 3A). We also measured a 20-fold decrease in TCT release by the TCT(-) strain compared to WT (Sup. Figure 3A). These findings confirmed previous data demonstrating the impact of AmpG efficiency on TCT release^11,15^.

**Figure 3:**
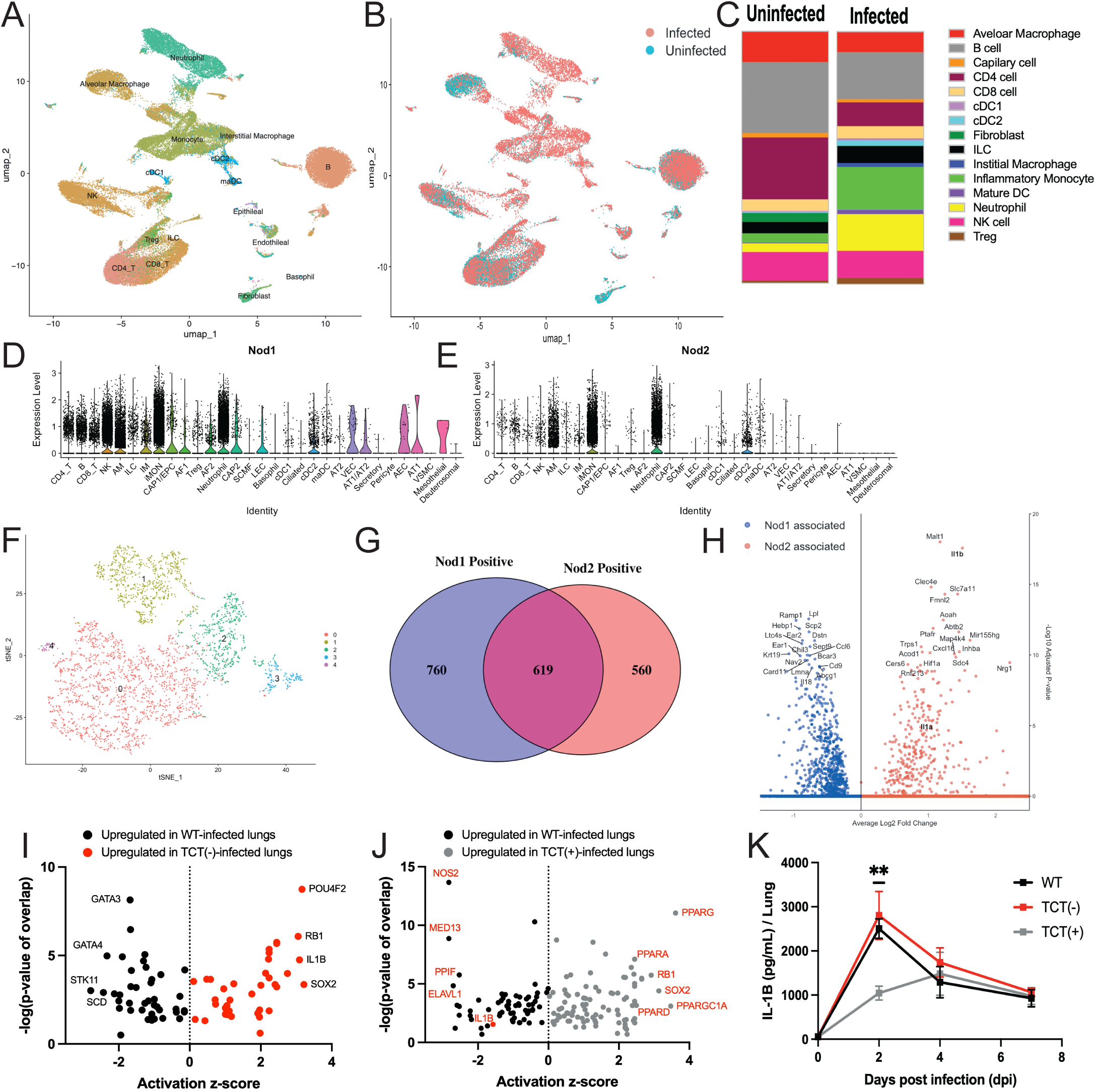
UMAP of (**A**) predicted cell types and (**B**) sample derived ID from single cell sequencing of PBS-sham challenged and *B. pertussis* infected C57/B6 mouse lungs at 4-dpi. (**C**) Comparison of relative abundance of cell types across infected and uninfected sample IDs. Cell type specific expression of (**D**) *nod1* and (**E**) *nod2* in the 4-dpi *B. pertussis* challenged mouse lung. (**F**) UMAP of subclustered alveolar macrophages (AM) from the single cell *B. pertussis* challenged mouse lung dataset. (**G**) Single cell analysis of 4-dpi alveolar macrophages assessed for NOD1 and NOD2 expression greater than 0.25 relative expression, grouping cells into NOD1+NOD2+, NOD1+NOD2-, NOD1-NOD2+, and NOD1-NOD2-. (**H**) Differential gene expression analysis comparing NOD1+NOD2- cells vs NOD1-NOD2+ cells derived from the 4-dpi single cell dataset. Upstream regulator analysis via the Ingenuity pathway analysis (IPA) pipeline comparing the 4-dpi lungs of BALB/c mice infected with (**I**) WT vs TCT(-) or (**J**) WT vs TCT(+), measured by Z-activation score for the relative upstream regulators annotated in the pipeline. (**K**) IL-1B lung protein levels assessed by ELISA for WT, TCT(-), and TCT(+) infected BALB/c mice at 2, 4, and 7-dpi.

To determine if AmpG efficiency could impact the structure of PGNs released extracellularly, we profiled muropeptides in the supernatant of WT, TCT(+) and TCT(-) cultures. We observed that the TCT(+) strain released, not only more TCT, but also greater amounts of almost all anhydrous-MurNAc containing muropeptides and saccharides, including GluNAc-anhMurNAc-tetrapeptide (TCT), GluNAc-anhMurNAc-tripeptide (GaM-Tri), GluNAc-anhMurNAc (GaM), anhMurNAc-tetrapeptide (aM-Tet), GluNAc-anhMurNAc-dipeptide (GaM-Di), and anhMurNAc-tripeptide (aM-Tri) (Figure 2E). Similarly, we observed greater enrichment of these PGN fragments in WT strain compared to TCT(-), confirming that *ampG* biases the release of anhydrous muropeptides (Figure 2E, Sup. Figure 3D) as they require access to the cytoplasm for processing to non-anhydrous forms ^24^. To determine the impact of efficient AmpG-mediated PGN recycling on extracellular PGN fragments efficiency on the release of non-TCT muropeptides we assesed fragments enriched in the TCT(-) strain than WT or TCT(+).

Excitingly, we identified multiple muropeptides which fit this profile. We observed greater release of GluNAc-MurNAc-Tripeptide (GM-Tri), MurNAc-Tripeptide (M-Tri), and MurNAc-Dipeptide (M-Di) in the TCT(-) strains conditioned media and decreased in the TCT(+) conditioned media (Figure 2E, F, Sup. Figure 3D). This suggests that loss of AmpG efficiency may prevent or reduce release of these structures. Excitingly, we hypothesize the enrichment of GM-Tri, M-Tri, and M-Di, all of which have been demonstrated to have potent NOD2 stimulatory capacity in other models, is driving NOD2 activation and the increased inflammation generated by the TCT(-) strain in mice. Interestingly, it was recently demonstrated that anhydrous MurNAc (aM) is a poor NOD2 agonist compared to non-reduced MurNAc which retains potent NOD2 stimulatory capacity when possessing a 2-4 amino acid stem peptide^25^.

Alternatively, we also observed enrichment of aM monosaccharide in the TCT(-) conditioned media and decreased amounts in the TCT(+) conditioned media (Sup. Figure 3C). Interestingly, this monosaccharide was the only PGN structure with anhydrous MurNAc that was also diminished in the TCT(+) conditioned media (Figure 3E). While the absence of the stem peptide should blunt any sensing by NOD1 or NOD2, recent advances in mammalian sensing of PGN suggest the possibility to sense this monosaccharide via an unidentified PGN sensor. MurNAc, unlike GluNAc, is a bacterial exclusive sugar, not produced by host cells, making this MAMP an interesting possible target for host PRRs but none has been identified to date^26,27^.

### NOD2 but not NOD1 drives inflammatory responses to *B. pertussis*

Improved AmpG mediated muropeptide transport reduced TCT release and increased the release of NOD2 agonist muropeptides (Figure 2E, F). Both NOD1 and NOD2 signal through the same adapters to drive similar NF-kB transcriptional responses and little is known about divergent outcomes of signaling via NOD1 or NOD2^28^. To determine why multiple bacterial species would dedicate such enormous metabolic output and risk immune detection we assessed outcomes of NOD1 and NOD2 signaling during *B. pertussis* infection. We hypothesized that host NOD2 drives a more potent inflammatory response. Thus, to determine the relative contributions of each NOD receptor we infected wild-type (WT), NOD1 (NOD1 KO) and NOD2 deficient (NOD2 KO) mice with *B. pertussis* and assessed the lung inflammatory pathology at 7-dpi.

Significantly immunopathology as measured by reduced bronchovascular bundle formation and alveolar consolidation was noted in NOD2 KO compared to WT mice (Figure 2G-I,K), suggesting NOD2 is a driver of host inflammation and immunopathology following *B. pertussis* infection. In contrast, NOD1 KO mice exhibited similar levels of inflammation relative to WT controls, though any potential increased inflammation in NOD1 KO mice may have been masked by the already substantial pathology observed in WT mice (Figure 2G-I, K). No differences in bacterial load were observed between WT, NOD1 KO and NOD2 KO mice (Figure 2J), indicating that the reduced inflammation in NOD2 KO mice was not due to differences in bacterial burden (Figure 2J).

To determine if the patterns of dampened neutrophil and NK cell recruitment observed by TCT release was dependent on NOD1 and/or NOD2 signaling, we assessed and compared the numbers of these immune cells in the lungs of WT, NOD1 KO, and NOD2 KO mice challenged with *B. pertussis*. Excitingly, we observed a similar pattern of reduced neutrophil and NK cell numbers in the lung in NOD2 KO mice, recapitulating the phenotype observed with greater TCT release suppressing the recruitment of these cells (Sup. Figure 5A, B). These findings further support the idea that NOD2 activating PGNs drive the recruitment of these key mediators of cellular immunity and that increased TCT release is a mechanism to avoid the release of these more pro-inflammatory muropeptide structures.

**Figure 4.**
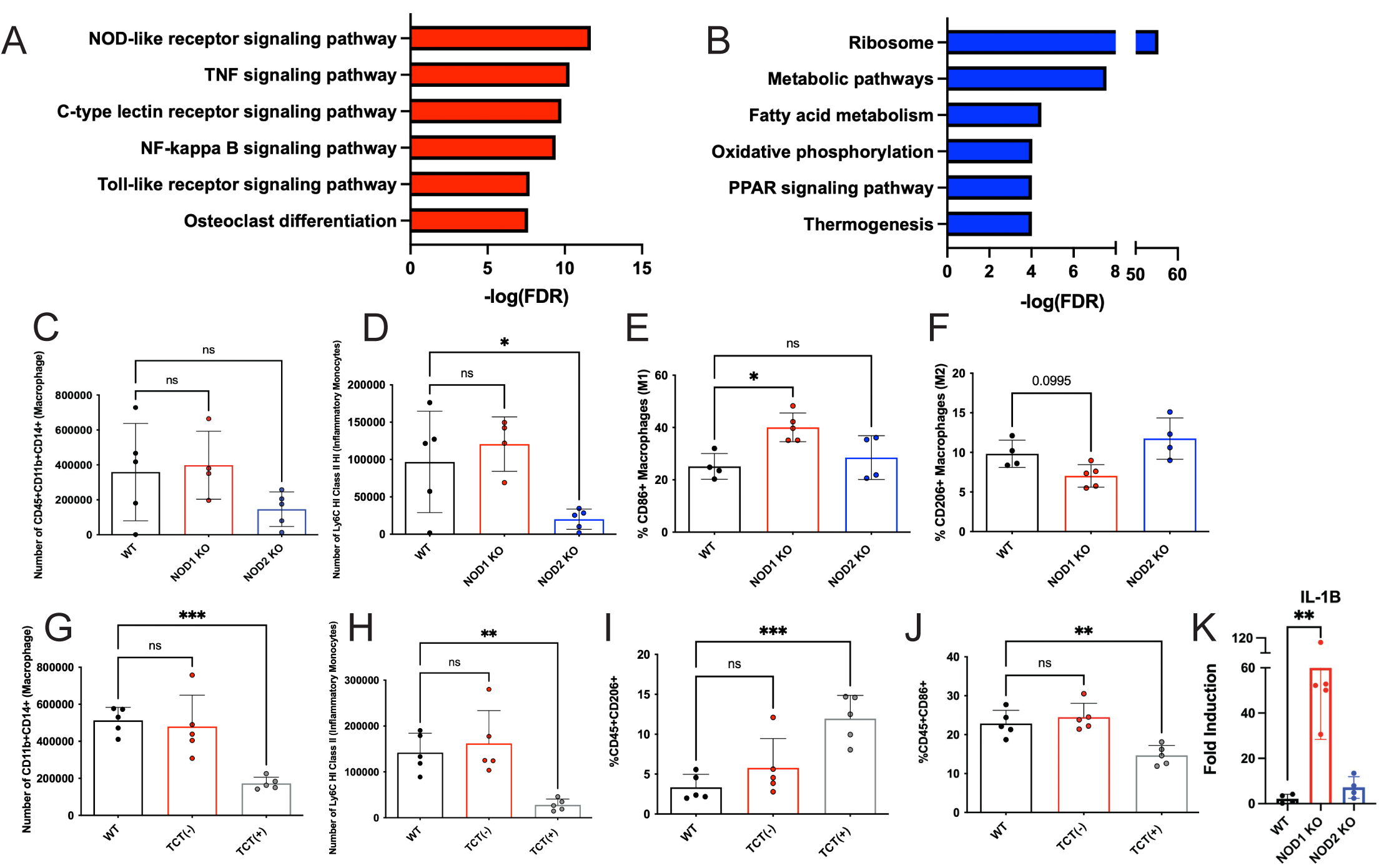
KEGG pathway enrichment analysis on differentially associated genes (FDR > 0.05) for (**A**) NOD1-NOD2+ and (**B**) NOD1+NOD2- alveolar macrophages, with infectious disease annotated pathways excluded from visualization. Number of (**C**) CD45+CD11b+CD14+ (Macrophages), (**D**) CD45+Ly6cHI ClassII+ (Inflammatory Monocytes), and percentage of (**E**) CD45+CD11b+CD14+CD86+ (M1 Macrophages) and (**F**) CD45+CD11b+CD14+CD206+ (M2 Macrophages) cells in the 4-dpi lungs of C57/B6 WT, NOD1 KO, and NOD2 KO mice challenged with WT parental strain of *B. pertussis*. Number of (**G**) CD45+CD11b+CD14+ (Macrophages), (**H**) CD45+Ly6cHI ClassII+ (Inflammatory Monocytes), and percentage of (**I**) CD45+CD86+ (M1-like) and (**J**) CD45+CD206+ (M2-like) cells in the 7-dpi lungs of C57/B6 WT, NOD1 KO, and NOD2 KO mice challenged with WT parental strain of *B. pertussis*. *i1lb* transcript levels in the 2-dpi lungs of *B. pertussis* challenged WT, NOD1 KO, and NOD2 KO mice, normalized to internal controls.

**Figure 5.**
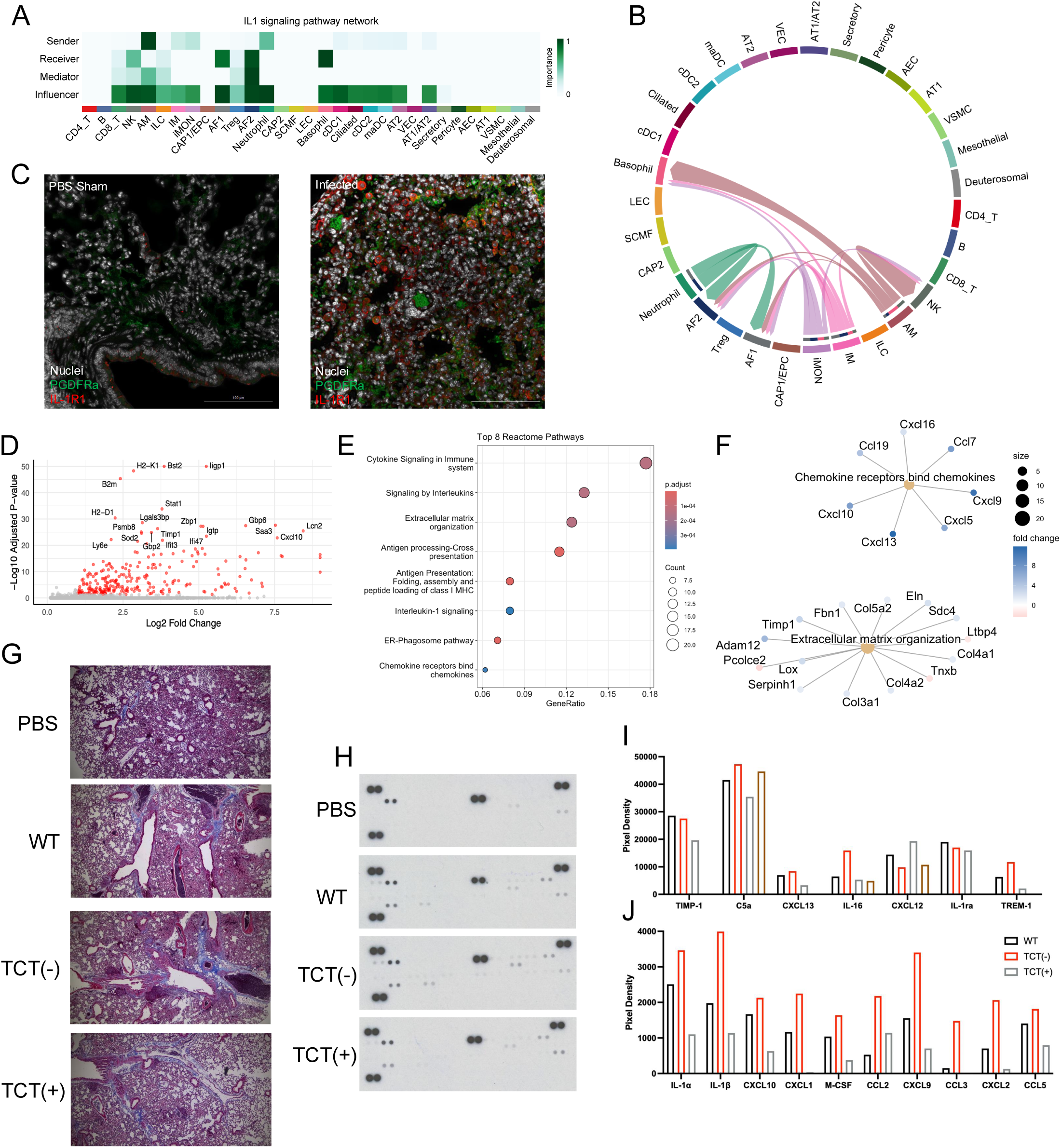
(**A** and **B**) CellChat analysis performed on single cell sequencing of PBS-sham challenged and *B. pertussis* infected C57/B6 mouse lungs at 4-dpi, identifying cell types predicted as involved in IL1 signaling. (**C**) Differential gene expression analysis performed on airway fibroblasts derived from infected (4-dpi) or uninfected mice then (**D**) Reactome pathway enrichment analysis performed, implicating (**E**) extracellular matrix reorganization and (**F**) chemokine pathways. (**G**) Representative 10x images of mason’s trichrome stain of 180-dpi lungs from mice challenged with WT, TCT(-), and TCT(+) strains of *B. pertussis*. (**H**) Proteome profiler array of pooled 7-dpi and PBS-sham lung lysates from WT, TCT(-), and TCT(+) challenged mice, quantified via ImageJ analysis (**I** and **J**).

### NOD2, but not NOD1, contributes to inflammatory cell programing during *B. pertussis* infection

We have identified divergent outcomes of NOD1 and NOD2 signaling during *B. pertussis* infection, with NOD2 but not NOD1 promoting inflammation (Figure 2). NOD1 and NOD2 signal through same adapters and activate the same transcription factors, so how these host receptors could be mediated different outcomes remained to be determined^28^. We hypothesized that differential expression of NOD1 and NOD2 across cell types and enrichment of NOD2 in inflammatory cell types would explain the previous findings. To investigate the contribution of NOD1 and NOD2 to inflammatory responses to *B. pertussis,* we performed single cell RNA sequencing on lungs from sham PBS-treated control and *B. pertussis* challenged (4-dpi) mouse lungs (Figure 3A, B). We used cell-type prediction algorithms to profile the cellular landscape during *B. pertussis* infection compared to the PBS sham challenged control, demonstrating an infiltration of inflammatory monocytes and neutrophils following infection (Figure 3A, C). Mapping NOD receptor expression across cell types revealed that NOD1 was broadly expressed, with abundant *Nod1* transcripts noted in nearly all cell types measured (Fig. 3C). But fitting with our expectation, NOD2 expression was more restricted, confined primarily to subsets of classically inflammatory cells like neutrophils, inflammatory monocytes (iMONs), as well as alveolar macrophages (AMs) and a subset of dendritic cells (cDC2s) (Figure 3D, E).

These findings support that NOD2 expression is associated with more pro-inflammatory cell types. However, to determine if NOD1 and NOD2 promote opposing outcomes within the same cell type we performed further analysis of AMs which were abundant in number and demonstrated robust and heterogenous expression of NOD1 and NOD2 (Figure 3C-E). Furthermore, AM localization to the alveoli and the ciliated cells of the epithelium suggest higher likelihood of direct interaction with *B. pertussis* and its effectors, e.g., TCT, implicating this cell type as critical early mediators to the differential outcomes observed with varying TCT release by *B. pertussis*.

To investigate and profile functional differences in AM populations, we performed subcluster analysis on these cells, identifying five transcriptionally distinct clusters of AMs, dubbed clusters 0-4 (Figure 3F). Two clusters (clusters 1 and 2) demonstrated significant upregulation of pro-inflammatory genes (Figure 3F, Sup. Figure 4A, B). We identified key markers for these clusters, and discovered cluster 1 was characterized by high expression *Ly6c2, Aoah, Nos2, Fmnl2, and Slamf7*, while cluster 2 was characterized by moderate expression of those genes in addition to high expression of *Msr1, Lrg1, Wfdc17, Ccl9, and Cxcl3* (Sup. Figure 5A). Nos2 has previously been linked to the effects of TCT *in vitro*, suggesting peptidoglycan-mediated responses could be driving the divergent AM cluster phenotypes^13^. We performed KEGG pathway analysis on each clusters’ differentially expressed genes to identify which pathways were upregulated in each AM cluster. NOD-like receptor signaling pathways was identified as the most enriched immune pathway in both clusters 1 and 2, further implicating the possible roles of NOD1 and NOD2 in shaping responses to *B. pertussis* infection (Sup. Figure 5C, D). We assessed NOD1 and NOD2 expression across these AM clusters and discovered that while NOD1 was ubiquitously expressed across each of the clusters, NOD2 expression was restricted primarily to more pro-inflammatory clusters 1 and 2, again highlighting NOD2 but not NOD1 in these responses (Sup. Figure 6B).

**Figure 6.**
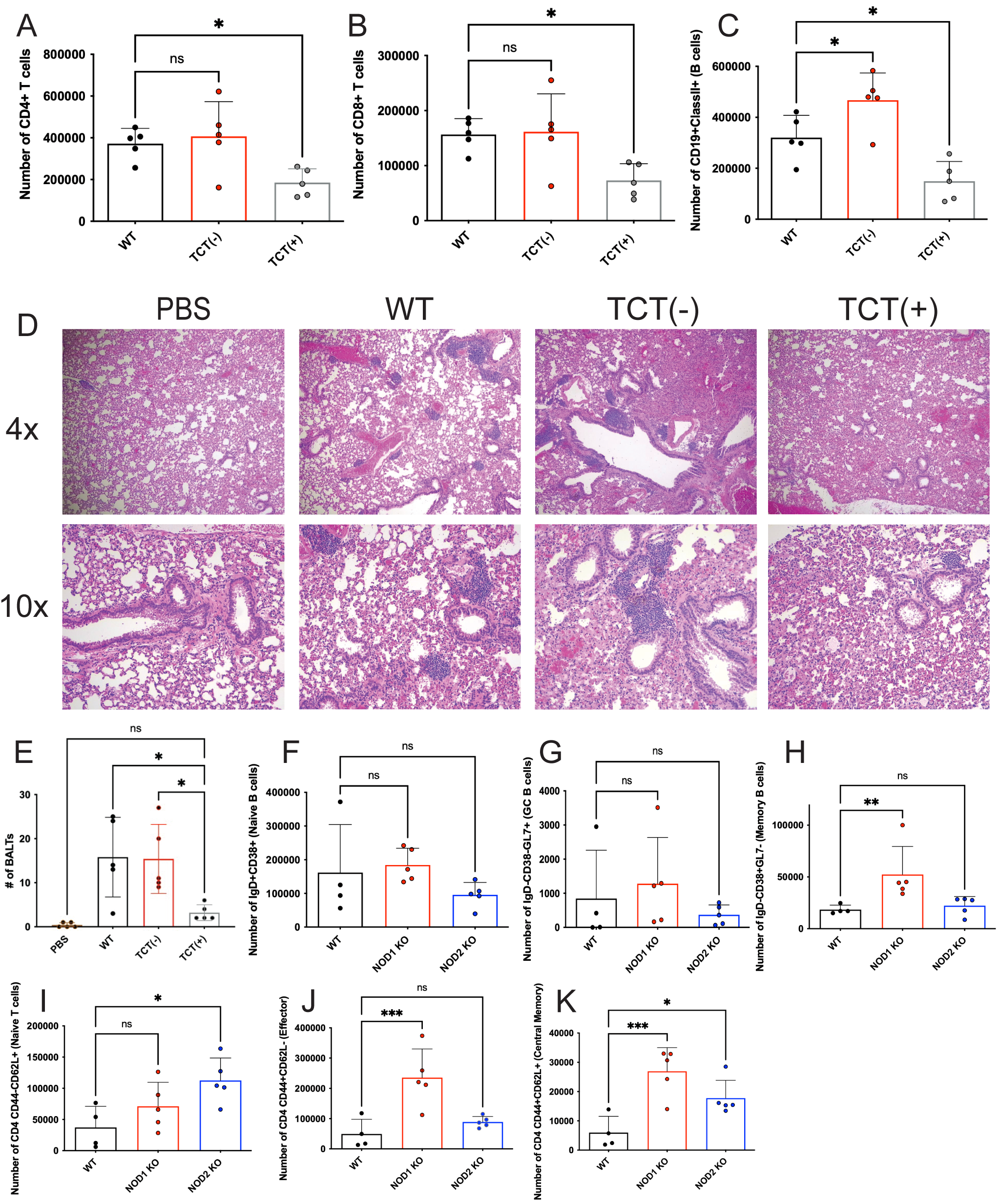
Number of (**A**) CD45+CD4+CD8- (CD4 T cells), (**B**) CD45+CD4-CD8+ (CD8 T cells), and (**C**) CD45+CD19+ClassII+ (B cells) in 7-dpi lungs of BALB/c mice challenged with *B. pertussis*. (**D**) Representative 4 and 10x images of H&E stains of PBS sham and *B. pertussis* strains WT, TCT(-), and TCT(+) 180-dpi BALB/c mouse lungs and **(E)** number of bronchi associated lymphoid tissue (BALT) in each 180-dpi lung sample. Number of (**F**) CD19+IgD+CD38+ (Naïve B cells), (**G**) CD19+IgD-CD38-GL7+ (Germinal Center B cells), (**H**) CD19+IgD-CD38+GL7- (Memory B cells), (**I**) TCRβ+CD4+CD44-CD62L+ (Naïve CD4 T cells), (**J**) TCRβ+CD4+CD44+CD62L- (Effector Memory CD4 T cells), (**K**) TCRβ+CD4+CD44+CD62L+ (Central Memory CD4 T cells) cells in 180-dpi lungs from C57/B6 WT, NOD1 KO, and NOD2 KO mice challenged with WT Tohama-1 derived parental strain of *B. pertussis*.

To delineate the roles of NOD1 and NOD2 in AMs, we stratified AMs based on NOD1+NOD2- and NOD1-NOD2+ expression profiles and performed differential gene expression analysis, excluding double-positive cells to isolate pathway-specific effects (Figure 3G). Distinct gene expression patterns emerged. Notably, NOD2+NOD1- AMs showed strong enrichment for *il1β* and *il1α* (Figure 3H). However, this link was not observed between NOD1 and IL-1 family cytokines in NOD1+NOD2- AMs (Figure 3H). We assessed IL-1α, and IL-1β expression in our subclusters of AMs and found that expression of these cytokines was heavily enriched in clusters 1 and 2, overlapping with expression of NOD2 but not NOD1 (Sup. Figure 4B). This further implicates NOD2 and not NOD1, as a driver of the pro-inflammatory program in AMs during *B. pertussis* infection. Surprisingly, while *il1a* and *il1b* were strongly associated with NOD2-expressing AMs, *il18*, another IL-1 signaling cytokine, was strongly associated with NOD1+NOD2- AMs (Figure 3H).

To determine if inefficient PGN transport and subsequent TCT release drives polarization of responses and immune outcomes, we investigated bulk-transcriptomic data comparing WT, TCT(+) and TCT(-) infected mouse lungs and performed Ingenuity Pathway Analysis (IPA) to predict upstream regulators driving differential gene expression. IPA identified IL-1β as amongst the top predicted upstream regulator of differences between WT and TCT(-) infected mice (Figure 3I). This further suggested that inefficient PGN recycling and subsequent TCT release dampens IL-1 driven inflammation and drives the pronounced inflammatory gene pattern observed between TCT(-) and WT infected mice (Figure 1H). Similarly, pro-inflammatory regulators (Nos2, IL-1β) were implicated amongst the top upstream regulators in the WT infected mice compared to the TCT(+) infected mice, validating the anti-inflammatory role of TCT and identifying macrophage polarization as a possible mechanism driving this phenomenon (Figure 3J).

Finally, to validate these transcriptomic predictions, we assessed the IL-1β protein levels in the lung over the course of *B. pertussis* infection by ELISA. At 2-dpi, we observed significantly higher IL-1β levels in mice infected with the WT strain compared to those infected with TCT(+) (Figure 3K), indicating that TCT release suppresses IL-1β production and confirming predictions of our upstream regulator analysis (Figure 3I, J). This data demonstrates that in response to *B. pertussis* PGN, NOD2 drives greater IL-1 production and that extracellular release of NOD1-agonist TCT by *B. pertussis* diverts away from this signaling pathway.

### TCT activation of NOD1 dampens M1 macrophage polarization and IL-1**β** production

Our data indicates that the inefficient PGN recycling of *B. pertussis* facilitates TCT release and dampens inflammatory responses to *B. pertussis* by activating polarizing NOD responses to decrease IL-1β production. However, the direct mechanisms by which NOD2, but not NOD1, contributed to IL-1β release remained unrefined (Figures 1-3). Single cell analysis of AMs suggested a possible skewing of M1 or M2 macrophage polarization as a possible mechanism (Figure 3F). Macrophages can be broadly characterized as having either an M1 or M2 phenotype^29^. M1 macrophages are associated with anti-microbial, and general inflammatory responses, while M2 macrophages are linked to tissue repair and immune regulation^30,31^. M1 polarization is characterized by IL-1β and iNOS (Nos2) production while M2 polarization is characterized by diminished expression of those pro-inflammatory mediators and instead greater expression of tissue homeostasis markers like Arg1, CD206 and IL-10^30,32–35^. Interestingly, we observed AM cluster populations 1 and 2, which were defined by their greater *Nos2* expression also expressed high levels of *Nod2* (Figure 3H). On the other hand, *Nod1* expression was abundant across all the subclustered populations of AMs (Figure 3H). *nos2* was significantly (FDR = 1.38e^-7^) associated with NOD2+NOD1- AMs, linking NOD2 and not NOD1 to M1 macrophages (Figure 3H).

Pathway enrichment analysis of *Nod2* associated genes in AMs identified the TNF signaling pathway as among the top enriched pathway, which is an effector response of M1 macrophages (Figure 4A)^36^. Furthermore, NOD-like receptor signaling pathway was implicated as the top differentially regulated pathway in NOD2+ AMs compared to NOD1+ AMs, suggesting NOD2 is greater driver of this signaling pathway during *B. pertussis* infection (Figure 4A). Conversely, pathway enrichment analysis of genes enriched in *Nod1+Nod2-* AMs revealed Ribosome, Metabolic pathways, Fatty acid metabolism, and PPAR (peroxisome proliferator-activated receptor) signaling pathway as the top enriched pathways in NOD1+ AMs (Figure 4B). PPAR signaling in macrophages is often a hallmark of M2-skewing^37^. Canonical M2 marker, CD206 (*mrc1*), was also identified as enriched in the NOD1+NOD2- AMs (FDR = 2.24e^-11^), further implicating NOD1and not NOD2 in promoting M2 macrophage differentiation.

To validate that NOD1 and NOD2 drive opposing M1/M2 skewing in response to *B. pertussis* challenge, we measured and profiled macrophage and monocytes in the lungs of NOD1 and NOD2 KO mice by flow cytometry. NOD2 KO mice exhibited significantly lower recruitment of inflammatory monocytes, and no difference in macrophages, compared to WT controls at 4-dpi (Figure 4C, D). To confirm our transcriptomic data and determine if NOD1 and NOD2 contribute to observed differences in macrophage polarization, we analyzed M1 and M2 macrophage subsets in the lungs of infected NOD1 and NOD2 KO mice. NOD1 KO mice displayed a significant increase in percent of M1-skewed macrophages (Figure 4E) and decreased (p=0.09) M2-like macrophages (%CD206+CD45+CD11b+CD14+) (Figure 4F). These data further support the hypothesis that TCT activation of NOD1 suppresses M1-like macrophage responses and promotes M2-like polarization during infection.

Upstream regulator analysis of bulk-RNA sequencing comparing WT and TCT(+) challenged mice, identified Nos2 as one of the least activated upstream regulators in the TCT(+) infected mice and PPAR as one of the most activated in TCT(+) infected mice, suggesting greater TCT release dampened iNOS production and promoted PPAR signaling (Figure 3J). Thus, our bulk RNA sequencing comparing mice infected with the different TCT strains suggests TCT inhibited M1-skewing and promoted M2-skewing. To determine if TCT release contributes to macrophage phenotype, we challenged mice with WT, TCT(-) and TCT(+) strains of *B. pertussis* and assessed the macrophage and monocyte numbers and M1/M2 polarization. TCT(+) challenge resulted in significantly reduced recruitment of macrophages (CD11b+CD14+) and inflammatory monocytes (Ly6cHI, Class II HI) in the lung at 7-dpi (Figure 4G, H), consistent with reduced inflammatory infiltrate observed in previous histological analysis (Figure 4G, H).

Next, we assessed how TCT influences macrophage/monocyte polarization during *B. pertussis* infection by comparing the percentage of M1 vs M2 macrophages in the lungs of mice challenged with WT, TCT(-) or TCT(+) strains. In keeping with our hypothesis, TCT(+)-infected mice exhibited a shift in immune polarization, with an increased frequency of M2-skewed immune cells (%CD45+CD206+) and a decrease in M1-skewed cells (%CD45+CD86+) relative to WT and TCT(-) challenge (Figure 4I, J). This shift is consistent with the model that greater TCT release by B*. pertussis* promotes pro-resolving M2-like macrophage response, typically associated with anti-inflammatory and tissue repair functions unlike pro-inflammatory M1 macrophages^29^. These results also fit with our earlier findings in which TCT(+)-infected mice demonstrated significantly less inflammatory pathology than WT-infected mice (Figure 1D).

As IL-1β production is a classical hallmark of M1 macrophage activation^32^, we measured *il1b* transcript levels in infected lungs. NOD1 KO mice showed a ∼60-fold increase compared to WT (Figure 4M), further supporting a role for NOD1 in limiting IL-1-mediated inflammation. These findings are consistent with reports from other inflammatory models in which NOD1 deficiency similarly exacerbated inflammation^38–40^. These data suggest TCT release promotes NOD1-dependent M2-skewing and avoids a NOD2-IL1 signaling axis. They also suggest that NOD1 may not just drive a weaker inflammatory response than NOD2 but rather contribute to a novel anti-inflammatory immune skewing, although the mechanism remains unknown.

### IL1 family cytokines reprogram airway fibroblasts, driving chemokine production and long-term lung remodeling following *B. pertussis* infection

Inefficient PGN recycling promotes release of TCT and dampens IL-1 signaling (Figures 1 and 4). To assess the potential benefits of this inefficient recycling to *B. pertussis* we assessed the role of IL-1 signaling in host immune responses to infection. CellChat analysis of single cell transcriptome data predicted neutrophils and AMs as the dominant producers (‘senders’) of *Il1α* and *Il1β* transcripts (Figure 5A, B), suggesting these immune cells drive the inflammatory response to *B. pertussis* (Figure 5A, B). CellChat analysis predicted that airway fibroblasts 1 and 2 (AF1, AF2) were the primary receivers of IL-1α and IL-1β with few other populations based on expression of IL-1 family receptors (Figure 5A, B). We also identified a putative signaling axis of IL-18-producing AMs signaling to IL18R+ basophils and NK cells (Figure 5B).

Airway fibroblasts are critical mesenchymal lineage cells which form and produce much of the cellular architecture of the lungs^41^. To validate these cells are presenting the IL1R1 receptor and thus have the capacity to respond to IL-1 during *B. pertussis* infection, we performed immunofluorescent staining of PBS sham and *B. pertussis* infected lungs, staining for IL1R1 and fibroblast marker, PDGFRa. In accordance with our single cell findings, we observed fibroblasts express IL1R1 receptor during *B. pertussis* infection (Figure 5C). To delineate how airway fibroblasts respond during *B. pertussis* infection, we performed differential gene expression analysis on fibroblasts derived from infected versus PBS-sham challenged lungs, followed by Reactome pathway enrichment analysis to identify genes and pathways enriched in airway fibroblasts during *B. pertussis* challenge (Figure 5D, E). Airway fibroblasts were found to express inflammation associated genes e.g., *Cxcl10, Zbp1* and *Saa3,* and IFN-inducible genes e.g., *Gbp6* and *Stat1* (Figure 5D). Furthermore, this analysis revealed robust upregulation of Interleukin-1 signaling pathway (Figure 5E), validating previous analysis suggesting airway fibroblasts are the main respondents to IL-1 cytokines during *B. pertussis* infection (Figure 5A, B). Additional enriched pathways included ‘Extracellular matrix (ECM) reorganization’ and ‘Chemokine receptors bind chemokines’ pathways (Figure 5D, E, F), suggesting that IL-1-activated airway fibroblasts may influence both the pronounced long-term pulmonary dysfunction and immune cell recruitment associated with *B. pertussis* infection (Figure 5D, E, F).

*\B. pertussis* infection can cause long-term pulmonary dysfunction, however, the mechanisms which mediate this are largely under-explored^42^. We hypothesized that IL1 family cytokine mediated activation of airway fibroblasts drives long-term collagen deposition and extracellular matrix reorganization following *B. pertussis* infection. As TCT limits IL-1 production we hypothesized that infection of mice with a TCT over-releasing strain would lead to reduced activation collagen deposition compared to challenge with WT *B. pertussis*. To test this, we performed long-term infection studies using WT, TCT(-) and TCT(+) *B. pertussis* strains. Lung tissue harvested at 6 months post-infection revealed increased collagen deposition in WT- and TCT(-)-infected mice, compared to TCT(+) (Figure 5G).

Differential gene expression analysis revealed significant upregulation of chemokines including *Ccl7, Cxcl9, Cxcl5, Cxcl13, Cxcl10, Ccl19*, and *Cxcl16* in airway fibroblasts following infection (Figure 5F). These findings are consistent with our earlier bulk RNA-seq data: *Cxcl13, Ccl19*, and *Cxcl5* were most highly expressed in TCT(-) infected lungs (Figure 1H), while *Cxcl19* and *Cxcl10* were downregulated in TCT(+) infections compared to WT (Figure 1I), mapping the fibroblast chemokine signature to the TCT-dependent chemokine expression patterns observed in the bulk RNA sequencing.

To validate bulk RNA sequencing data proteome profiling was performed on lung lysates from WT, TCT(+) and TCT(-) infected mice (Figure 4H). TCT(-) infection induced more IL-1α and IL-1β than WT or TCT(+) challenge (Figure 5I, J). Broadly, we observed higher chemokine levels in the TCT(-) group compared to the WT and TCT(+) infected mice (Figure 5I, J). Notably, protein levels of Cxcl13, Cxcl19, and Cxcl10 were highest in TCT(-) infected lungs and lowest in TCT(+) infected lungs (Figure 5I, J), mirroring gene expression trends and reinforcing the conclusion that airway fibroblasts are robust producers of chemokines and shape immune responses to *B. pertussis* following IL-1-mediated activation.

### TCT release skews NOD signaling to dampen IL-1**β** mediated adaptive cell recruitment and immune memory formation

We have established that TCT release dampens the expression and production of multiple chemokines (Fig. 1, 5), including CCL19, CXCL9, CXCL10 and CXCL13. CXCL13 is essential for recruiting B cells to germinal centers and inflamed tissues, while CXCL9, CXCL10 and CCL19 mediate T cell recruitment to sites of inflammation^43–46^. Based on this data, we hypothesized that TCT-mediated suppression of chemokine responses would impair adaptive immune cell infiltration into the lung and formation of germinal centers. To determine if TCT influences adaptive immune cell recruitment, we performed flow cytometry on 7-dpi lungs of mice infected with WT, TCT(-), or TCT(+) strains of *B. pertussis*. TCT release inversely correlated with recruitment of adaptive immune cells (Figure 6A-C). TCT(+) infected mice demonstrated decreased numbers of CD4, and CD8 T cells, and B cells in the lungs at 7-dpi compared WT infected mice, suggesting TCT release may be a mechanism to dampen recruitment of these cells (Figure 6A-C). Furthermore, TCT(-) infected mice, had significantly greater B cells in the lungs compared to WT infected mice, further implicating inefficient PGN recycling and TCT release in blocking recruitment and formation of adaptive immunity (Figure 6A-C).

To assess the consequences of TCT release on long-term immune responses we infected mice with WT, TCT(-) or TCT(+) *B. pertussis* strains and assessed lungs immunopathology at 6-months post-infection by H&E stain (Figure 6D). We observed formation and persistence of long-lived lymphoid structures in mice infected with *B. pertussis* compared to PBS challenged controls (Figure 6D). At 6 months post-infection, immune bronchus-associated lymphoid tissue (iBALT) structures were markedly reduced in the lungs of TCT(+) infected mice, suggesting that TCT suppresses the formation of long-lived lymphoid structures, limiting adaptive immune memory (Figure 6D, E).

As TCT is a NOD1 exclusive agonist, and our data suggests that NOD2 drives a stronger IL1-mediated induction of adaptive responses from airway fibroblasts than NOD1 and that NOD1 signaling during *B. pertussis* infection results in anti-inflammatory outcomes. We hypothesized that inefficient PGN recycling and TCT release and stimulation of NOD1 is beneficial for *B. pertussis* as it inhibits the formation of adaptive immunity, which may be bolstered in the presence of NOD2 activating PGN structures. To investigate this, we assessed the role of NOD1 and NOD2 in the formation of long-term immunity to *B. pertussis*. WT, NOD1, and NOD2 KO mice were infected with *B. pertussis* and lung-resident memory lymphocytes profiled at six months post-infection. Naïve B cell frequencies and germinal center B cell numbers were similar across groups (Figure 6F, G), but NOD1 KO mice exhibited significantly elevated lung resident memory B cells (IgD-CD38+GL7-) compared to WT mice (Figure 6G). Similarly, both central (CD44+CD62L+) and effector (CD44+CD62L-) memory CD4 T cells were increased in NOD1 KO mice (Figure 6J, K), while naïve CD4+ T cell numbers remained unchanged (Figure 6H). These findings suggest that NOD1 signaling restricts adaptive memory formation, suggesting that inefficient PGN recycling and TCT release may be a mechanism to dampen or prevent long-term immune memory. In NOD2 KO mice, we observed a modest increase in central memory CD4+ T cells, accompanied by an increase in naïve T cells, suggesting an expanded T cell pool rather than enhanced memory (Figure 6I, K) and highlighting the potential impact of TCT biasing of NOD responses.

We hypothesize that TCT inhibits M1 macrophage polarization to dampen IL-1-mediated recruitment of immune cells by airway fibroblasts. To determine if inhibition of IL-1 signaling could impede immune cell recruitment and bacterial clearance, we challenged WT and IL1R1 KO mice with *B. pertussis* and evaluated inflammatory pathology, bacterial burden, and immune cell profiles. IL1R1 KO mice exhibited impaired bacterial clearance (Figure 7A), highlighting the essential role of IL-1 in pathogen control. Notably, despite higher bacterial burdens, IL1R1 KO mice exhibited significantly reduced pulmonary inflammation (Figure 7B), confirming IL-1’s role as a central inflammatory mediator, and supporting our hypothesis that IL1 signaling is a mediator of the NOD2-dependent *B. pertussis* inflammation we observed (Fig. 2E-G).

**Figure 7.**
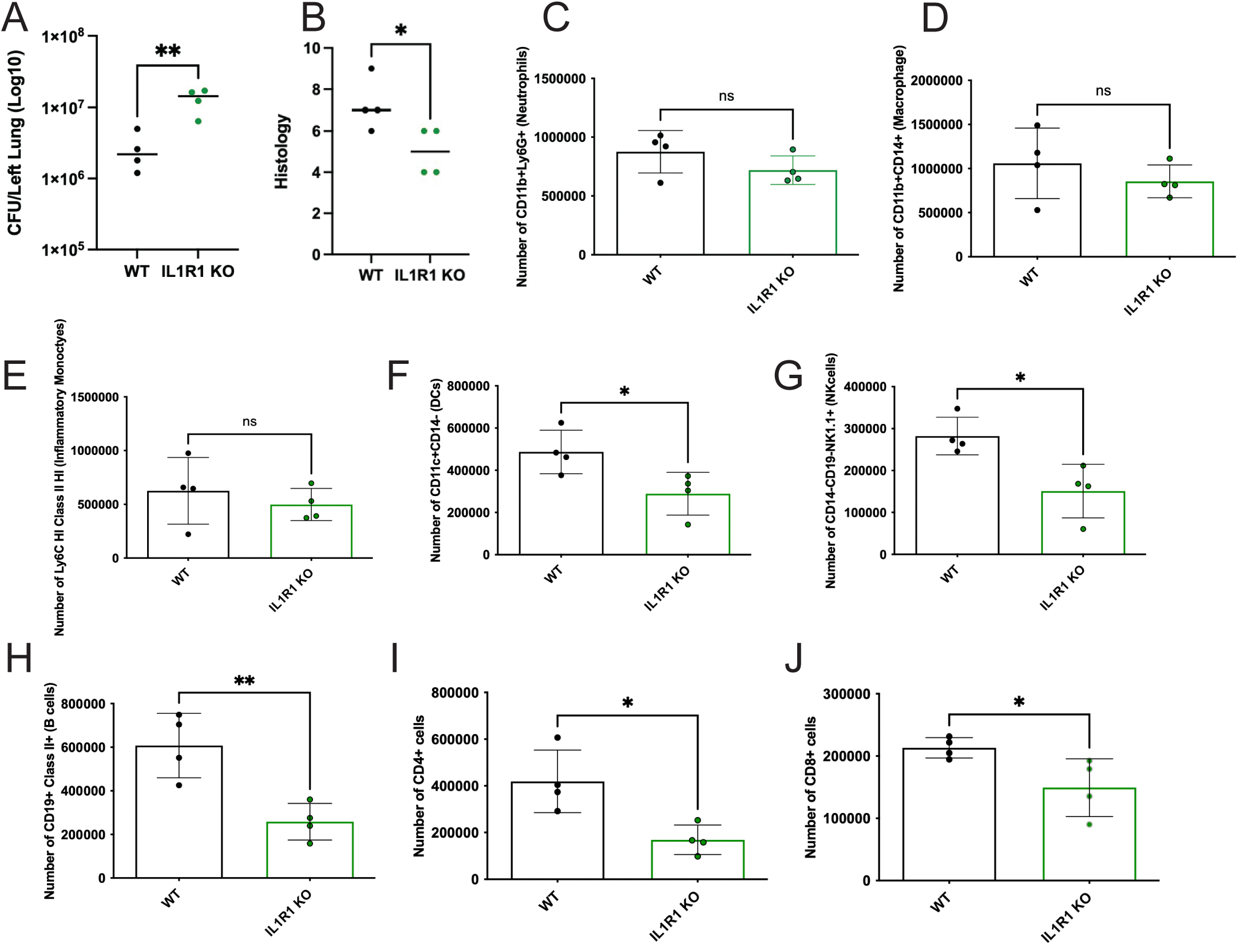
(**L**) Bacterial load and (**M**) inflammatory histology score of H&E stained 7-dpi lungs from C57/B6 WT and IL1R1 KO mice intranasally challenged with 2x10^6^ CFU. Number of (**N**) CD45+CD11b+Ly6g+ (Neutrophils), (**O**) CD45+CD11b+CD14+ (Macrophages), (**P**) CD45+Ly6cHI ClassII+ (Inflammatory Monocytes), and (**Q**) CD45+CD11c+CD14- (Dendritic cells), (**R**) CD45+CD19+ClassII+ (B cells), (**S**) CD45+CD4+CD8- (CD4+ cells), and (**T**) CD45+CD4-CD8+ (CD8+ cells) cells in the 7-dpi lungs of C57/B6 WT and IL1R1 KO mice challenged with WT strain of *B. pertussis*.

Surprisingly, innate immune cell recruitment (e.g., neutrophils, macrophages, and monocytes) was similar between WT and IL1R1 KO mice (Figure 7C-E) despite IL1R1 KO mice demonstrating reduced bacterial control and clearance (Figure 7A), supporting the model that early recruitment of neutrophils to the lung does not contribute control of infection^47^. Yet strikingly, recruitment of lymphocytes like NK, B and T cells was significantly diminished in IL1R1 KO mice compared to WT, indicating that IL-1 signaling is necessary for adaptive immune cell infiltration into the lung (Figure 7G-J). Furthermore, a modest but significant decrease in dendritic cells was also observed in infected IL1R1 KO compared to WT controls, suggesting the bridging of the innate to adaptive response may be further hindered (Figure 7F). Taken together, these findings support our hypothesis that TCT promotes *B. pertussis* immune evasion by skewing NOD1-driven responses to suppress a NOD2-mediated activation of AMs and release of IL-1 cytokines to drive the activation of IL1R1 expressing airway fibroblasts. Activation of these fibroblasts leads to enhanced chemokine expression, recruitment of adaptive immune cells and formation of germinal centers. This limits inflammation and compromises the development of protective immunity.

## DISCUSSION

This study reveals a novel immune evasion strategy employed by *Bordetella pertussis*, selective release of a peptidoglycan fragment, TCT, which activates NOD1 while avoiding NOD2 engagement. This strategy parallels the way many Gram-negative bacteria modify their lipid A to modulate TLR4 engagement. We propose that *B. pertussis* has evolved a similar level of structural specificity in its PGN release to manipulate intracellular PRR signaling and evade immunity.

*B. pertussis* release of TCT during the turnover of saccular PGN is essential for bacterial growth and division. *B. pertussis* release of NOD1-activating TCT diminishes the release of NOD2 activating muropeptides. This strategy biases the host response toward NOD1 signaling, which we demonstrate results in less inflammation than activation of NOD2. This represents a previously unrecognized form of immune evasion based on selective release of structurally distinct cell wall fragments and highlights an unappreciated difference in disease outcomes following engagement of NOD1 or NOD2.

While both NOD1 and NOD2 signal through NF-kB, our data reveal distinct functional consequences *in vivo*^28^. NOD2 promotes pulmonary inflammation and facilitates immune memory, but NOD1 suppresses inflammation and restrains the development of adaptive immunity. These data fit with recent in-silico work suggesting anhydrous peptidoglycan structures have anti-inflammatory roles, by blocking recognition of NOD2 and biasing towards NOD1^48^. These findings are supported by single-cell RNA-sequencing analyses demonstrating that NOD1 is broadly expressed across many cell types in the infected mouse lung, but NOD2 expression is restricted to inflammatory cells, including neutrophils, monocytes, alveolar macrophages and cDC2’s. This restricted expression of NOD2 may contribute to its inflammatory potency. Additionally, NOD2’s dual caspase recruitment (CARD) domains and its ability to engage mitochondrial antiviral signaling protein (MAVS) and type I IFN pathways, may further distinguish its activity from NOD1^49–51^. Type I and III IFNs have been implicated in exacerbating pertussis disease while contributing little to nothing towards bacterial control and clearance, therefore NOD2 signaling contributing to these pathways is detrimental to the host^52^.

We identify a key downstream mechanism by which NOD2 signaling contributes to pulmonary pathology. Specifically, NOD2-driven IL1 production activates IL1R1+ airway fibroblasts, which in turn upregulate genes associated with extracellular matrix remodeling, collagen deposition and chemokine secretion. This fibroblast activation promotes immune cell infiltration and iBALT formation but also contributes to long-term tissue remodeling and fibrosis. In contrast, TCT limits IL1 production by avoiding NOD2 signaling, reducing fibroblast activation, chemokine secretion and immune infiltration. These findings connect PGN structure to fibroblast activity and immune organization in the lung.

Interestingly, while IL1 was dispensable for neutrophil and monocyte recruitment, it was critical for B and CD4+ T cell infiltration and pathogen clearance. In IL1R1-deficient mice, we observed increased bacterial burden and impaired adaptive immunity, despite intact innate responses. Previous work by others in the field have linked IL-1β to bacterial clearance^53^. These results highlight the central role of the NOD2-IL1-fibroblast axis in orchestrating effective long-term immunity to *B. pertussis*. By skewing this axis, TCT promotes bacterial survival and immune evasion.

TCT release impaired the formation of iBALT and limited memory B and T cell development. These data suggest that early inhibition of IL1 production and fibroblast activation by TCT has long lasting consequences for adaptive immunity. Importantly, these findings explain how paradoxically *B. pertussis* may evade long-term immune surveillance and establish persistent infection by selectively exposing immune polarizing PGNs. Assessments of released PGN structures in previously and currently circulating clinical strains may provide further evidence of an evolutionary pressure on the availability and structure of muropeptides released by *pertussis*^54^. Further, these data have implications for other extracellular PGN producing infections such as *Neisseria gonorrhoeae* and *Borrelia burgdorferi*^9,55^ for which the development of effective vaccines has been a persistent challenge.

These data also have direct implications for pertussis vaccine development. We suggest that selecting or engineering muropeptides that robustly stimulate NOD2 may improve immune priming, while vaccine strains lacking TCT could avoid the immunosuppressive effects we describe. Notably, the live attenuated *B. pertussis* vaccine candidate BPZE1 was engineered to lack TCT release^15,56^. Our data provide critical mechanistic support and biological justification for this approach, demonstrating that TCT release suppresses IL-1 production, restrains adaptive immune cell recruitment, and limits long-term immunity. Blocking TCT release, as in BPZE1, or adjuvanting with NOD2 activating molecules may therefore enhance immunogenicity and protection of future pertussis vaccines.

## METHODS

### Mice and ethics statement

All animal procedures were approved by the Institutional Animal Care and Use Committee at the University of Maryland School of Medicine (protocol 00000108). Adult (6–10-week-old) BALB/c and C57BL/6, NOD1 - /- NOD2 -/-, SLC46A2 -/-, and IL1R1 -/- mice were bred and housed under specific pathogen-free conditions.

### Bacterial strains and infections

*B. pertussis* strains wild-type BC36, a streptomycin-resistant Tohama I derivative, isogenic mutants TCT(-), or TCT(+) were cultured on Bordet-Gengou agar supplemented with 10% defibrinated sheep blood or in Stainer-Scholte (SS) broth. Growth curves were generated by incubating strains of *B. pertussis* in 96-well plates, in 250 uL volume of SS broth, at starting ODs of 0.1, 0.05, and 0.01. Plate was incubated at 37C, shaking, and the OD was read every 10 minutes by the Cerillo Optical Density Microplate Reader. *B. pertussis* TCT mutant strains were generously supplied by William Goldman (University of North Carolina at Chapel Hill). TCT(+) strain was generated by expression of the *E. coli* AmpG in BC36, resulting in 24-fold increase in TCT release^11^. TCT(-) was generated by deletion of native *B. pertussis* AmpG and insertion of a kanamycin resistance gene at the site, resulting in a 50-fold decrease in TCT release^57^. Adult mice were anesthetized and inoculated intranasally with 2 × 10⁶ CFU of *B. pertussis* in 50 µL PBS^58^. Lungs were harvested at 2, 4, 7, 14, 180 days post-infection for bacterial burden analysis by serial dilution and plating of homogenates, tissue sectioning and staining, RNA extraction, protein preparation, single cell preparation of lung-derived cells, and for flow cytometry analysis.

### Histopathology and collagen staining

Lungs were fixed in 10% formalin, embedded in paraffin, and sectioned at 5 µm thickness. Slides were stained with hematoxylin and eosin (H&E) or mason’s trichrome stain. A blinded veterinary pathologist scored sections for peribronchial and perivascular inflammation, alveolitis, and airway epithelial damage^59^. Bronchus associated lymphoid tissue (BALT) formation were tallied in the 180-dpi H&E stained lungs.

Images were acquired using a Zeiss Axio Imager.M2 microscope equipped with an Axiocam 506 color camera and a Zeiss EC Plan-Neofluar 40x Pol objective (420363-9901). Image acquisition and processing (stitching, unsharp masking, and export) were performed using ZEN software version 3.10 (Zeiss).

### Liquid chromatography and mass spectrometry analysis

*B. pertussis* WT, TCT(-), and TCT(+) strains were grown to mid-log in S.S. media and the conditioned media was isolated from these cultures via centrifugation and filtering. 1.0 mL samples were taken from each culture and adjusted to 0.1% trifluoracetic acid before filtering through 0.22 µm filter prior to analysis. Samples were loaded on Acquity UPLC BEH C18 column 2.1 x 50 mm (Waters) using a S11 Dionex UHPLC. Samples were eluted over a 6-minute linear gradient of 0% to 50% acetonitrile (0.1% formic acid) in water (0.1% formic acid) before analysis on Q-Exactive Orbitrap (Thermo Fisher Scientific). All data were processed and analyzed on a Thermo Xcalibur Qual Browser.

### Immunofluorescence staining and microscopy

To process lung tissue, we paraffin-embedded and sectioned (0.5 µm), then placed on glass slides. Samples were prepared for sequential staining through deparaffinization in consecutive xylene washes, followed by rehydration in decreasing concentrations of ethanol ranging from 100% to 50%, antigen retrieval, and blocking steps^60^. The mounted tissue samples were incubated with the according primary and secondary antibody, following previously published methods^61^. Immunofluorescence images of the stained lung specimens were captured using an Olympus CSU W1 Spinning Disk Confocal Microscopy System. Image analysis for quantification of fluorescence signal was conducted through the Imaris 3D Analysis Software (v10).

### Flow cytometry analysis

The left lobe of the lung was collected, and single cell suspensions were generated using ACK lysis and collagenase digestion following Cell STAR Protocols method for single-cell suspension preparation from murine organs, lung section^62^. After counting, cells were washed once with 1X PBS and stained for viability using Ghost Dye™ UV450 Viability dye (Tonbo Biosciences) diluted 1:1000 in 1X PBS for 10 minutes on ice. After viability staining, cells were washed once with 1X PBS containing BSA, sodium azide, and EDTA (FACS buffer). Fc receptors were blocked for in a solution containing 100µg/mL each of ChromPure™ mouse, rat, and hamster IgGs (Jackson ImmunoResearch Laboratories Inc.) in FACS buffer for 15 minutes on ice. Surface staining was performed next in FACS buffer for 15 minutes on ice. After surface staining, cells were washed twice with FACS buffer. Samples were analyzed on the Cytek Aurora 4, and all gating and compensation was generated using Flow Jo 10.8.1 (BD Biosciences).

### RNA isolation and processing

Lung tissue was harvested at 2, 4, or 7-dpi for RNA isolation by the Trizol-choloroform method. Lung tissue was homogenized using the Omni Bead Beader (Omni, Inc), phase separated using chloroform and precipitated using isopropanol. RNA was then quantified and converted to cDNA (iScript cDNA Synthesis Kit). Quantitative real-time PCR (qPCR) was performed using the BIORAD CFX96 real-time PCR instrument. Gene expression was calculated as fold change relative to PBS-inoculated control animals using the 2^−ΔΔCT threshold cycle method and normalized to Hypoxanthine phosphoribosyltransferase (HPRT) (internal housekeeping gene).

### Bulk RNA sequencing

At 4-dpi, lungs were harvested from WT BALB/c mice (n = 3/group) infected with either WT, TCT(-), or TCT(+) strains of *B. pertussis*. Total RNA was extracted from homogenized lungs using Trizol Reagent (Invitrogen) according to the manufacturer’s protocol. RNA integrity was confirmed using a Bioanalyzer (Agilent), and sequenced on an Illumina NextSeq 1000. FastQC and RSeQC were used to assess read and alignment quality. Reads were aligned with the *Mus musculus* GRCm39 reference genome. Differential gene expression was analyzed using DESeq2, with significance defined as adjusted p-value (FDR) < 0.05. QIAGEN proprietary Ingenuity pathway analysis (IPA) sub-package Upstream Regulator was performed on genes identified as differentially expressed between groups (WT vs TCT(-) and WT vs TCT(+)) and ranked by Z-activation score.

### Single-Cell RNA sequencing

Single-cell suspensions were prepared from lung tissues of duplicate PBS-sham treated and Tohama-1 *B. pertussis* infected WT C57BL/6 mice at 4-dpi. Single cell preparation followed the published and validated Cell STAR Protocols method for single-cell suspension preparation from murine organs, lung section^62^.

Following red blood cell lysis, cell viability was assessed by trypan blue staining, viable cells were enriched by (Miltenyi Biotec) Dead Cell Removal Kit and loaded onto the 10x Genomics Chromium Controller for barcoding and library construction. Sequencing was performed on an Illumina NovaSeq platform. Data were processed using Cell Ranger and analyzed using the Seurat v5 package in R. Quality control cutoffs of >1000 and <10,000 features and <20% mitochondrial RNA. Data were normalized via the standard Seurat workflow (NormalizeData, FindVariableFeatures, ScaleData) and individual Seurat processed samples were integrated via the Harmony method. LungMap Mouse CellCards Multi-Study CellRef 1.0 Atlas single cell dataset was utilized as a reference with known cell types to generate predictions for cell types in our single cell dataset. Clustering was performed via the Seurat package workflow (ScaleData, RunPCA, FindNeighbors, FindClusters, RunUMAP(dims = 1:30), RunTSNE(dims = 1:30). Clusters were annotated by the hallmark variable features determined by the software. DeSeq2 was performed to determine the differential gene expression between alveolar macrophages assessed as either NOD1 positive (0.25 expression level cutoff), NOD2 negative (0.25 expression level cutoff) and NOD2 positive, NOD1 negative. CellChat analysis performed on single cell dataset. Reactome pathway enrichment analysis performed on differential expressed genes identified (Wilcox) from PBS-sham and infection derived airway fibroblasts. Data visualized by Seurat (DimPlot, Vlnplot, DotPlot) and ggplot2 (Volcano, HeatMap) R packages.

### Lung protein preparation and quantification

At 2, 4 and 7-dpi, lungs were harvested from WT BALB/c mice (n = 5/group) infected with either WT, TCT(-), or TCT(+) strains of *B. pertussis*. Lungs were homogenized in PBS with protease inhibitors and treated with Triton X-100 (1%). Samples were freeze-thawed at -80C and then centrifuged to remove cellular debris.

7DPI lung samples (n = 5) were pooled for each group (PBS, WT, TCT(-), and TCT(+)) and processed by bio-techne Proteome Profiler Array, Mouse Cytokine Array Panel A (Catalog # ARY006) protocol. Resulting array was quantified by ImageJ analysis. 2, 4, and 7-dpi lung samples (n = 5) were assessed by bio-techne Mouse IL-1 beta/IL-1F2 ELISA Kit (Catalog # MLB00C) protocol.

### Reporter cell assays for NOD1 and NOD2 activation

HEK293-derived reporter cells expressing murine or human NOD1 and NOD2 with an NF-kB inducible SEAP reporter (InvivoGen) were stimulated with *B. pertussis* tracheal cytotoxin, TCT (generously provided by William Goldman, UNC), live *B. pertussis* strains (MOI of 400), and the conditioned media from the bacteria grown in Stainer-Scholte (SS) media. *B. pertussis* strains were grown to log phase, and the spent media was collected at OD 0.8. Spent media was centrifuged and filtered to remove bacteria and large fragments. Supernatants were analyzed for reporter activity using colorimetric (NOD1 and NOD2 reporter cells; InvivoGen) detection kits (InvivoGen).

## Acknowledgments

We acknowledge Vincent Bruno for providing assistance with QIAGEN Ingenuity Pathway Analysis. Funding generously provided by R01 AI141372 N.H.C., NIH/NIAID grant #30049751 A.J.S., R01 AI167947-01 C.S., T32 AI095190 D.M.R..

## Author contributions

D.M.R. and C.S. conceived and designed the project. D.M.R., E.M.H., B.C, R.E.H., S.C., C.S., K.P., T.B.C., and A.J.S, performed experiments. D.M.R., E.M.H,. and B.C analyzed results. R.B. and N.S. designed and provided mice strains. C.L.G., K.C.S., N.H.C., M.C.G., and W.E.G. provided reagents and consultation. The paper was written by D.M.R. and C.S..

## Declaration of interests

The authors declare no other competing interests.

**Supplemental Figure 1.** (**A**) Growth curves of WT, TCT(-), and TCT(+) strains grown in SS broth. Number of (**B**) CD45+CD11b+Ly6g+ (Neutrophils), (**C**) CD45+CD11b+CD14+ (Macrophages), (**D**) CD45+Ly6cHI ClassII+ (Inflammatory Monocytes), and (**E**) CD45+CD11c+CD14- (Dendritic cells), in the 4-dpi lungs of BALB/c mice challenged with WT, TCT(-), or TCT(+) strains of *B. pertussis*.

**Supplemental Figure 2.** (**A**) Murine or (**B**) human NOD1 response to purified TCT and C12-iE-DAP control, and (**C**) Murine or (**D**) human NOD2 response to purified TCT and L18-MDP control, measured by HEK293 reporter cells with an NF-kB based SEAP reporter, assessed at 655nm.

**Supplemental Figure 3.** Relative abundance of TCT in the conditioned media of WT, TCT(-), and TCT(+) strains of *B. pertussis*, measured by LC-MS and visualized in the LC runoff time (**A**) and peak area of the curve (**B**). (**C**) Relative abundance of anhydrous MurNAc (aM) in the conditioned media of WT, TCT(-), and TCT(+) strains of *B. pertussis*, measured by LC-MS, visualized in peak area. (**D**) Relative abundance of PGN fragments found in the conditioned media of WT and TCT(-) strains of *B. pertussis*, measured by LC-MS, visualized in ion count.

**Supplemental Figure 4.** Number of (**A**) CD45+CD11b+Ly6g+ (Neutrophils) and (**B**) CD45+NK1.1+ (NK) cells in the 4-dpi lungs of C57/B6 WT, NOD1 KO, and NOD2 KO mice challenged with WT parental strain of *B. pertussis*.

**Supplemental Figure 5.** (**A**) Markers of subclustered alveolar macrophages, and (**B**) cluster specific expression for *nod1*, *nod2*, *il1b*, and *il1a* from the single cell *B. pertussis* challenged mouse lung dataset. KEGG pathway analysis of differentially expressed genes for alveolar macrophage clusters (**C**) 1 and (**D**) 2 compared to other subclusters of alveolar macrophages in the *B. pertussis* challenged single cell 4-dpi C57/B6 murine lungs.

