## Supplementary figures and images for "*B. pertussis* tracheal cytotoxin biases NOD signaling to suppress IL-1 mediated inflammation and evade adaptive immunity"

### Supplemental Figures

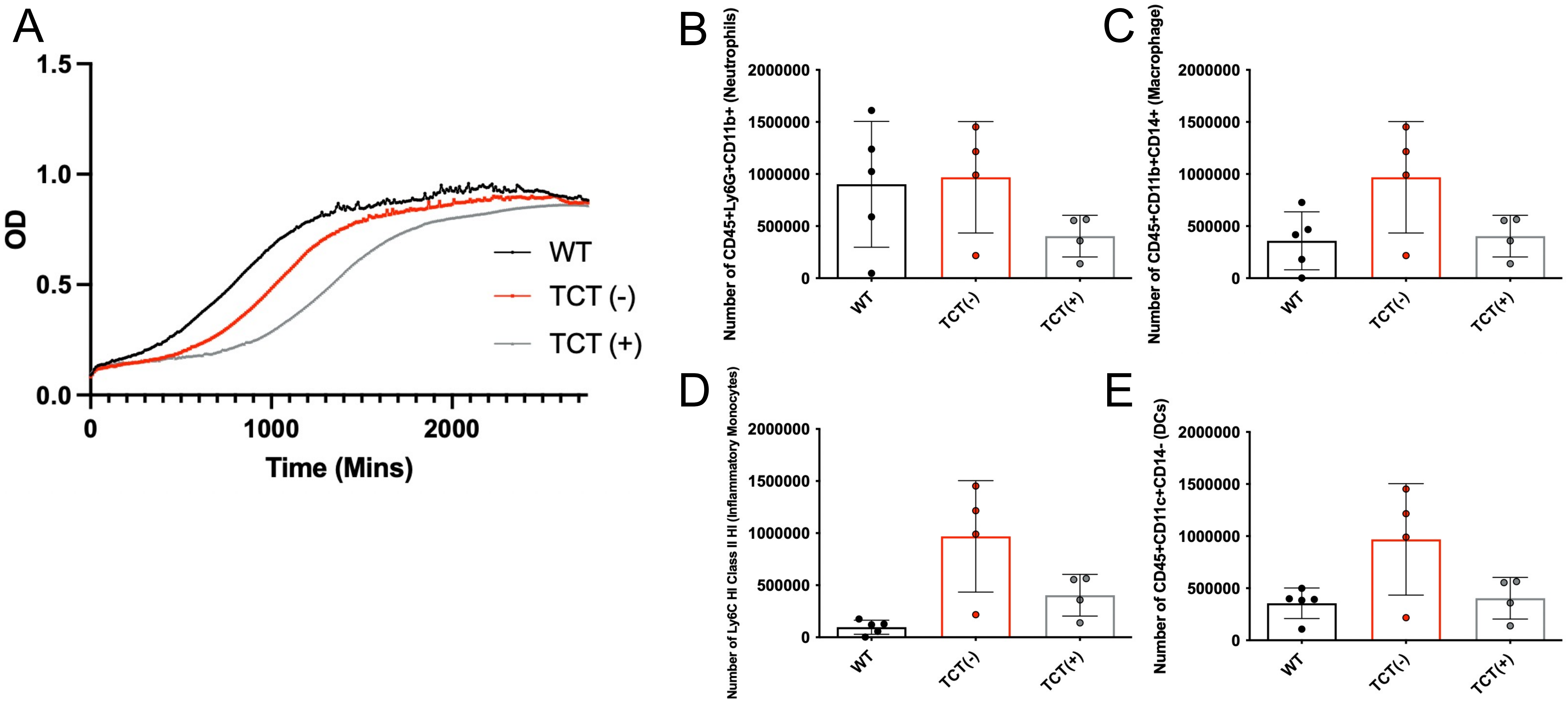

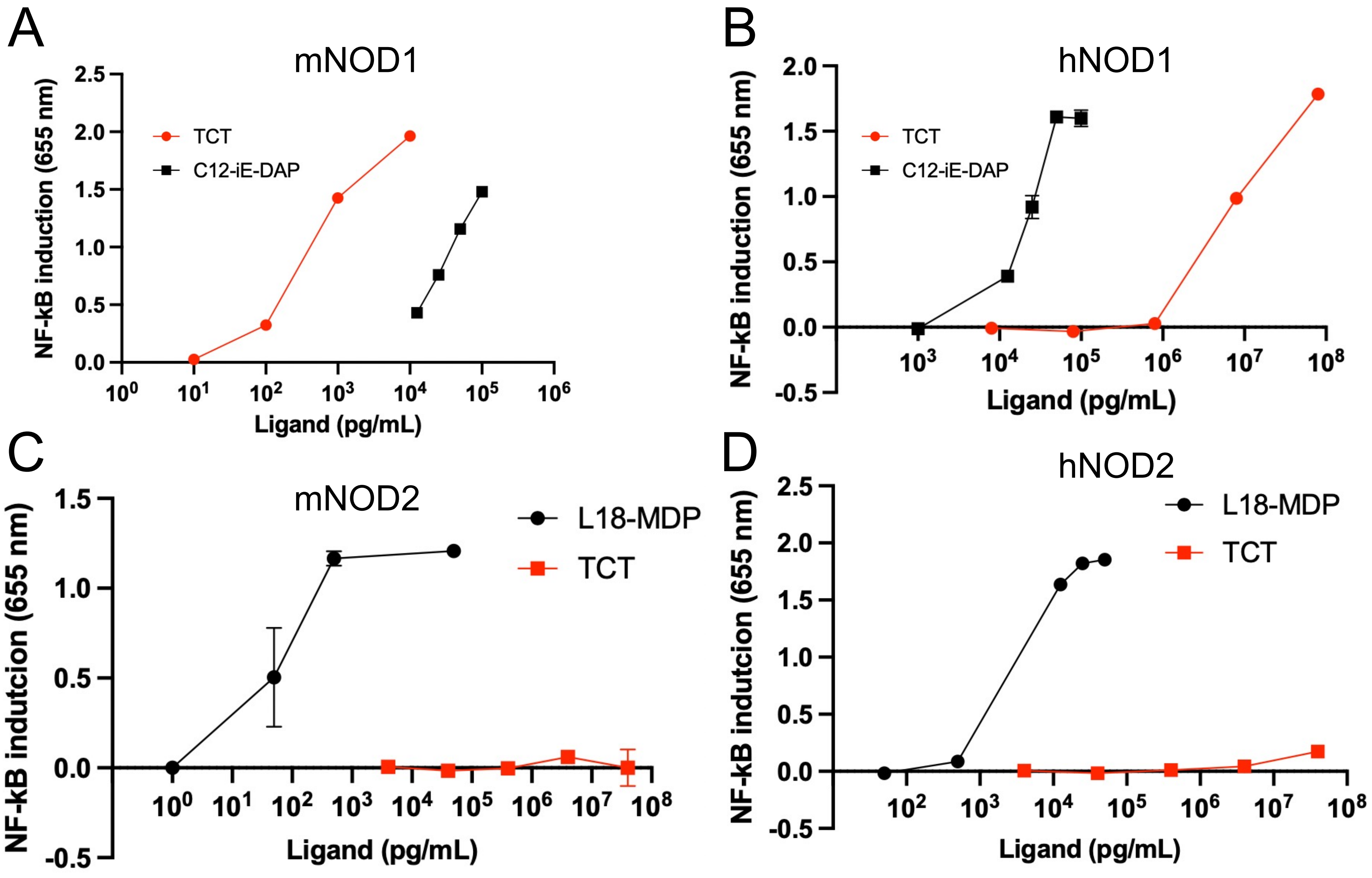

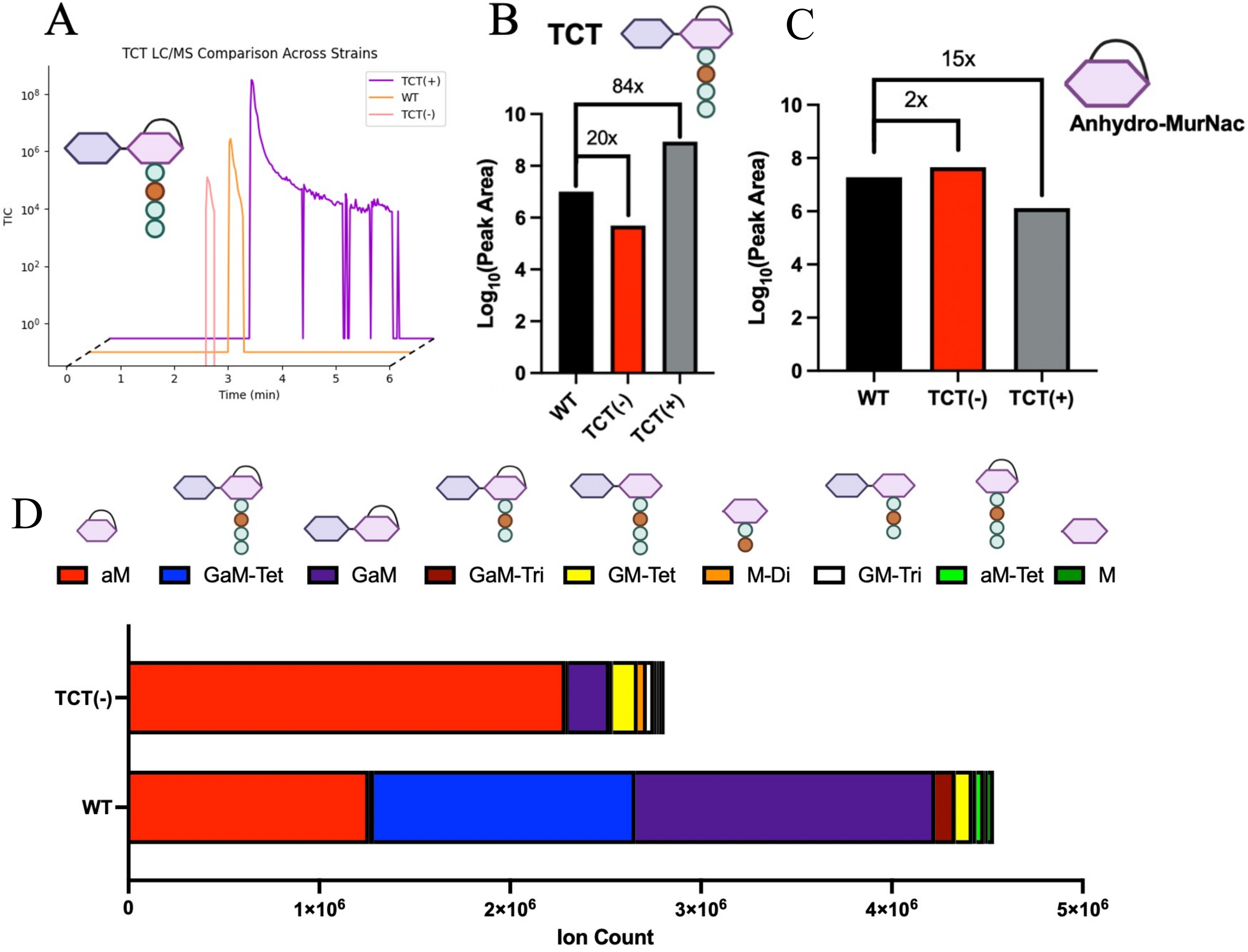

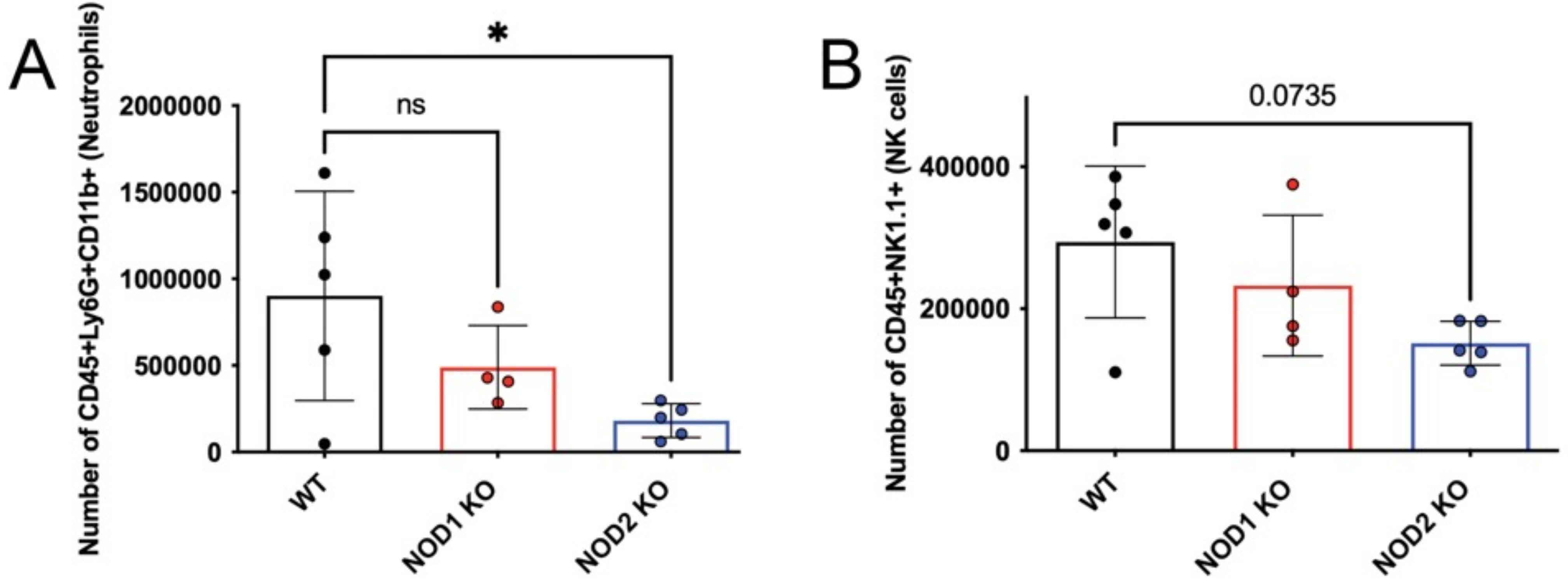

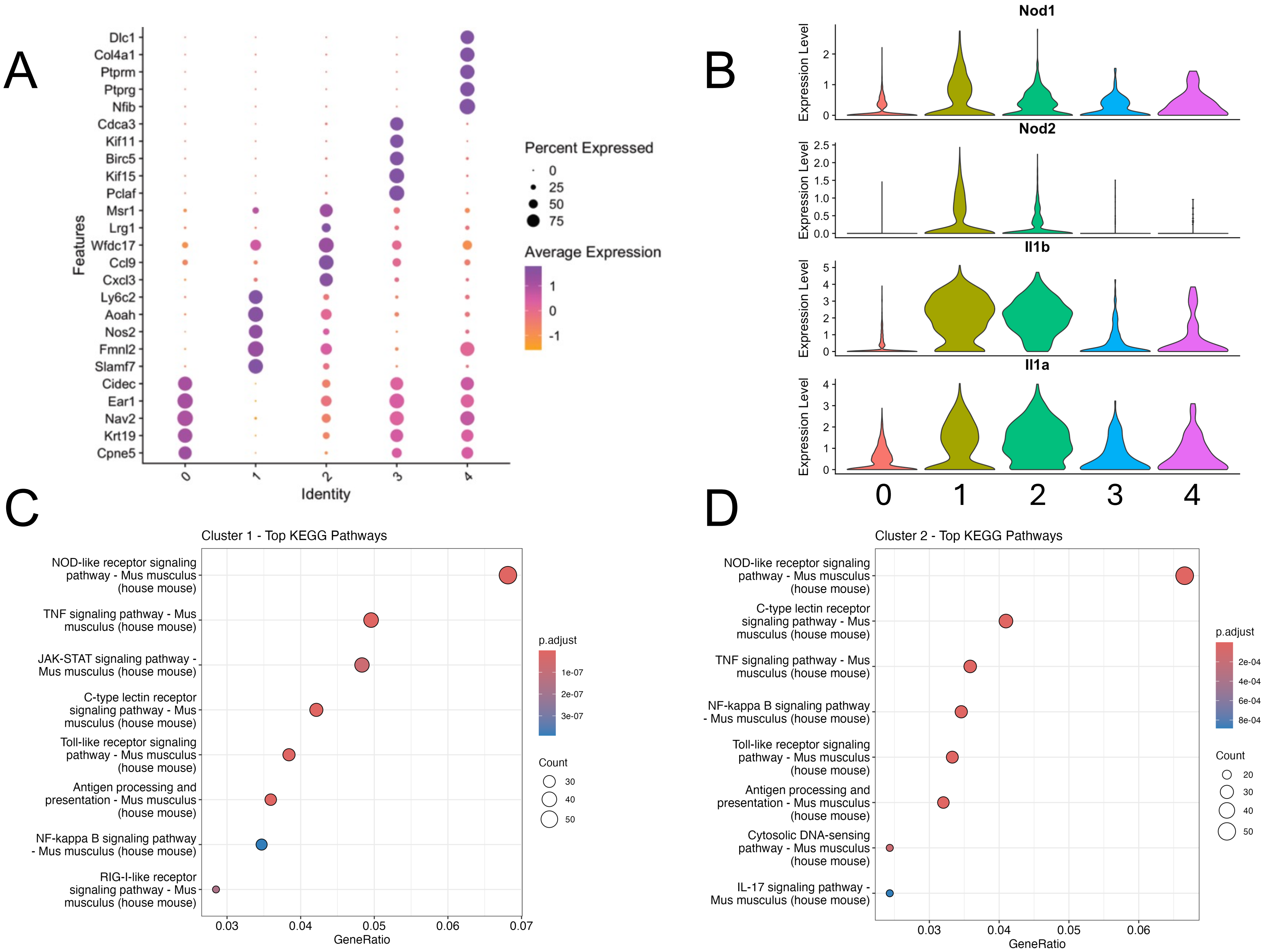
